# Multi-omic responses to acute exercise in abdominal subcutaneous adipose tissue of sedentary adults: findings from MoTrPAC

**DOI:** 10.64898/2026.03.05.702363

**Authors:** Cheehoon Ahn, Christopher A. Jin, Katie L. Whytock, Gina M. Many, Tyler J. Sagendorf, James A. Sanford, Zhenxin Hou, Mark R. Viggars, Jia Nie, Sara E. Espinoza, Nicolas Musi, Yifei Sun, Maria F. Pino, Patrick Hart, Dan H. Katz, Hasmik Keshishian, Gregory R. Smith, Scott Trappe, Natalie M. Clark, Sue C. Bodine, Laurie J. Goodyear, Karyn A. Esser, Christopher B. Newgard, Bryan C. Bergman, Joshua N. Adkins, Martin J. Walsh, Lauren M. Sparks, MoTrPAC Study Group

**Affiliations:** Translational Research Institute, AdventHealth, Orlando, FL, USA; Department of Genetics, School of Medicine, Stanford University, Stanford, California, USA; Biological Sciences Division, Pacific Northwest National Laboratory, Richland, WA, USA; Department of Biochemistry, Emory University School of Medicine, Atlanta, GA, USA; Department of Physiology and Aging, University of Florida, Gainesville, FL, USA; Department of Medicine, Cedars-Sinai Medical Center, Los Angeles, CA, USA; Icahn School of Medicine at Mount Sinai, New York, NY, USA; Proteomics Platform, Broad Institute of MIT and Harvard, Cambridge, MA, USA; Department of Medicine, Stanford University School of Medicine, Stanford, CA, USA; Human Performance Laboratory, Ball State University, Muncie, IN, USA; Aging and Metabolism Research Program, Oklahoma Medical Research Foundation, Oklahoma City, OK, USA; Section on Integrative Physiology and Metabolism, Joslin Diabetes Center, Harvard Medical School, Boston, MA, USA; Duke Molecular Physiology Institute and Sarah W. Stedman Nutrition and Metabolism Center, Duke University Medical Center, Durham, NC, USA; Division of Endocrinology, Diabetes, and Metabolism, University of Colorado Anschutz Medical Campus, Aurora, CO, USA

## Abstract

Exercise induces widespread health benefits across multiple tissues, yet the acute molecular responses in human adipose tissue remain poorly defined. The Molecular Transducers of Physical Activity Consortium (MoTrPAC) profiled temporal molecular changes in abdominal subcutaneous adipose tissue (ASAT) following a single bout of exercise. Healthy sedentary adults were randomized to endurance (EE), resistance (RE), or control (CON) groups. ASAT biopsies were collected pre-exercise and at 45min, 4hr, and 24hr post-exercise, followed by transcriptomic, proteomic, phosphoproteomic, and metabolomic analyses. EE and RE elicited distinct, time-resolved molecular programs involving angiogenesis, extracellular matrix remodeling, mitochondrial metabolism, substrate utilization, and circadian regulation. Phosphoproteomics revealed acute changes in cytoskeletal and branched-chain amino acid metabolism proteins associated with glycemic control. Temporal metabolomic shifts were cell-type-specific. Finally, we identified candidate adipose-derived exerkines with predicted endocrine actions. This multi-omic map of acute ASAT responses offers insight into adipose-specific mechanisms by which exercise promotes metabolic health.

## INTRODUCTION

The cardiometabolic health benefits of regular exercise are well-established^1,2^, yet the molecular mechanisms that drive these effects are incompletely understood. Many of these benefits—such as enhanced whole-body and tissue-specific insulin sensitivity and reduced blood pressure^3–5^—are initiated by single bouts of exercise, even in individuals who exercise regularly^6–8^. It is widely accepted that the cumulative effects of these repeated, episodic molecular responses to acute exercise bouts underlie the long-term health benefits of training^9^. As such, mapping the molecular landscape of acute exercise responses offers a promising strategy to uncover ‘how’ exercise promotes cardiometabolic health.

While much of the current and historical focus has been on skeletal muscle and the cardiovascular system, adipose tissue is emerging as a key target of exercise adaptations. Abdominal subcutaneous adipose tissue (ASAT) responds dynamically to both acute exercise^10–12^ and chronic training^10,13–17^, contributing to improved cardiometabolic outcomes. Moreover, similar to other tissues, ASAT may also mediate the systemic effects of exercise^18^ by serving as a rich source of numerous secreted factors—often referred to as exerkines—that act across tissues^19^. However, the physiological adaptations and molecular-level responses in human ASAT, particularly in the context of temporal and modality-specific dynamics of acute exercise, remains a significant gap.

The Molecular Transducers of Physical Activity Consortium (MoTrPAC) was launched to address this knowledge gap by systematically mapping the molecular effects of physical activity. In this study, which leveraged pre-COVID MoTrPAC human cohort data, we profiled temporal, multi-omic responses in ASAT following a single bout of either endurance exercise (EE) or resistance exercise (RE)—both of which represent clinically relevant exercise prescriptions at an unprecedented scale. Here, we reveal temporal-, modality-, and cell-type-specific molecular responses in the ASAT transcriptome, proteome, phosphoproteome, and metabolome following acute exercise. We examine the clinical relevance of these molecular signatures and, through *in silico* modeling, explore the potential endocrine role of ASAT-derived exerkines. Together, our findings provide new insights into the mechanisms underlying early ASAT-specific molecular responses to exercise and identify candidate mediators of long-term metabolic benefits.

## RESULTS

### Participant characteristics and multi-omic sex differences

173 healthy, sedentary individuals were enrolled and randomized into one of three groups: an endurance exercise group (EE, N=63), a resistance exercise group (RE, N=73), or a non-exercise control group (CON, N=37) (Figure 1A). The cohort was predominantly female (72%), with 46 women in EE, 49 in RE, and 31 in CON, with an average age of 41±15 years and a BMI of 27.1±4.0 kg/m² (Table 1). Participants in EE completed a 40-minute bout of endurance exercise at 65% VO2peak, while those in RE performed an 8-exercise resistance circuit at volitional fatigue (∼10RM), and the CON group rested in the supine position for 40 minutes (Figure 1A). Abdominal subcutaneous adipose tissue (ASAT) biopsy samples were collected before and at post 45min, 4hr, and 24hr (Figure 1A). To mitigate the confounding effects that may stem from a prior biopsy on the same side (e.g., biopsy-induced inflammatory response), participants were randomized into either “Early” (n=43; Pre-Exercise and post 45min), “Middle” (n=80; Pre-Exercise and post 4hr), or “Late” (n=50; Pre-Exercise and post 24hr) groups that underwent a total of two ASAT biopsies (one on either side of the abdomen) that included a pre-exercise (all groups) and one post-exercise timepoint (Figure 1A). Collected ASAT samples were subjected to four omic analyses: transcriptomics, global proteomics, phosphoproteomics, and metabolomics (Table S1).

**Figure 1.**
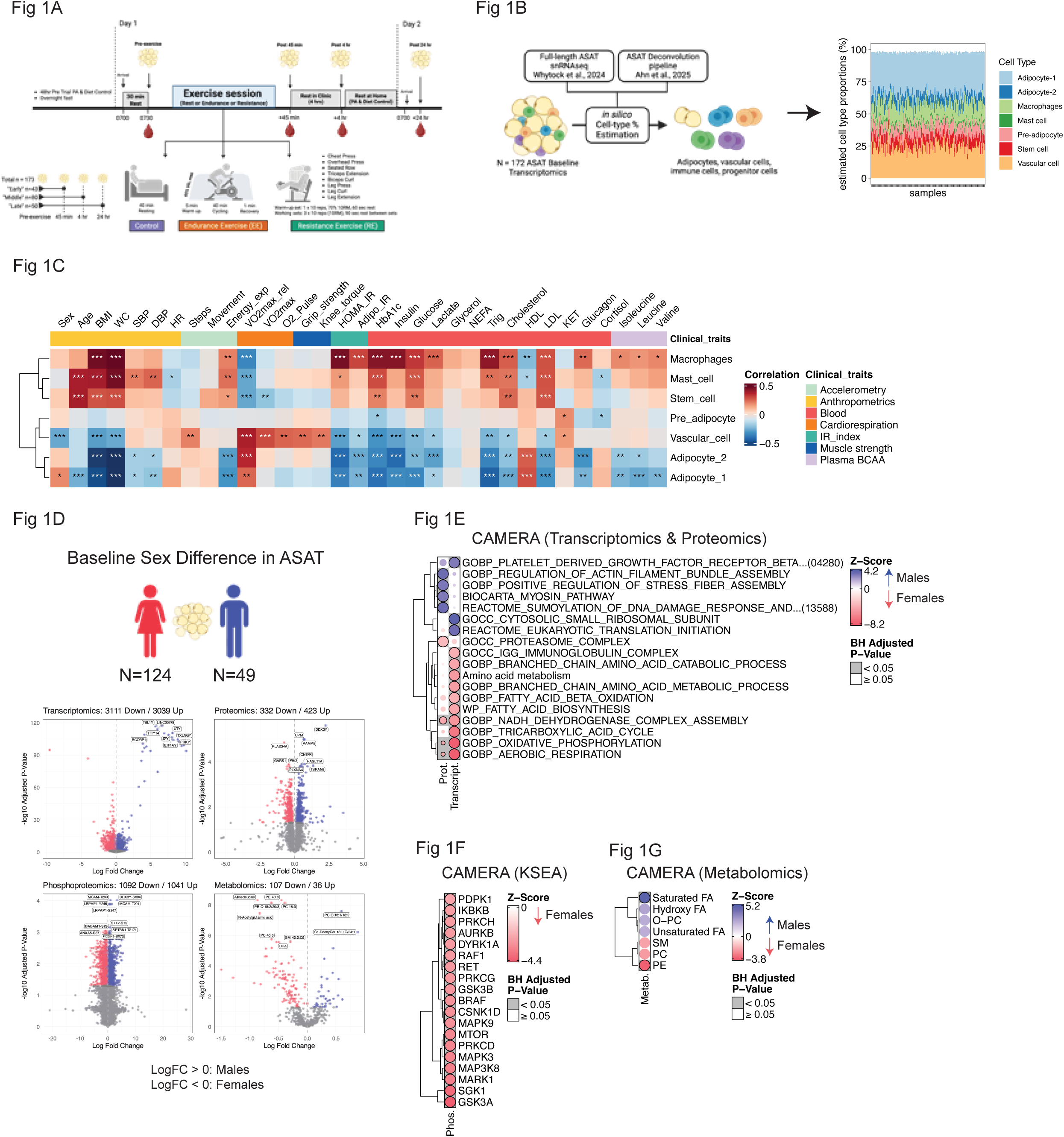
Study design and baseline characteristics. A) Overview of the study design. B) Workflow of pre-exercise cell type deconvolution of ASAT. C) Heatmap showing correlation between estimated cell type proportions and clinical/subclinical variables. “p<0.05, **p<0.01, ***p<0.001. D) Volcano plots showing sex-based differences (male vs. female) in transcriptomics, proteomics, phosphoproteomics, and metabolomics. E) CAMERA-PR results highlighting pathways enriched in males vs. females. Positive z-scores indicate male-biased enrichment. For Kinase substrate enrichment analysis (KSEA), no kinases were significantly upregulated in the males.

**Table 1.** Pre-exercise participant characteristics. BMI, Body Mass Index; HR, Heart Rate; SBP, Systolic Blood Pressure; DBP, Diastolic Blood Pressure; eGFR, estimated glomerular filtration rate; HDL, High-Density Cholesterol; LDL, Low-Density Cholesterol; HOMA-IR, Homeostatic Model Assessment for Insulin Resistance; NEFA, Non-esterified fatty acids.

| Characteristic | Overall<br>N = 173<sup>1</sup> | CON<br>N = 37<sup>1</sup> | EE<br>N = 63<sup>1</sup> | RE<br>N = 73<sup>1</sup> |
| --- | --- | --- | --- | --- |
| Age (yrs) | 41 (15) | 42 (15) | 41 (14) | 40 (15) |
| Height (cm) | 168 (9) | 167 (8) | 167 (9) | 168 (10) |
| Weight (kg) | 76 (13) | 73 (13) | 76 (13) | 77 (14) |
| BMI (kg/m<sup>2</sup>) | 26.9 (4.0) | 26.2 (3.9) | 26.9 (3.9) | 27.2 (4.0) |
| Waist Circumference (cm) | 92 (11) | 89 (13) | 92 (10) | 93 (12) |
| Systolic BP (mmHg) | 116 (12) | 114 (11) | 117 (11) | 116 (12) |
| Diastolic BP (mmHg) | 72 (8) | 72 (8) | 73 (9) | 72 (8) |
| Hemoglobin A1c (%) | 5.30 (0.30) | 5.29 (0.21) | 5.28 (0.32) | 5.31 (0.32) |
| eGFR (mL/min/1.73 m<sup>2</sup>) | 101 (20) | 99 (19) | 103 (18) | 101 (22) |
| Total Cholesterol (mg/dL) | 192 (40) | 197 (44) | 192 (40) | 189 (39) |
| HDL Cholesterol (mg/dL) | 58 (15) | 62 (17) | 57 (14) | 56 (15) |
| LDL Cholesterol (mg/dL) | 117 (34) | 119 (38) | 116 (34) | 116 (32) |
| HOMA-IR | 2.21 (1.52) | 1.99 (2.07) | 2.07 (1.18) | 2.44 (1.46) |
| NEFA (mmol/L) | 0.54 (0.19) | 0.57 (0.19) | 0.56 (0.21) | 0.51 (0.17) |
| **Cardiopulmonary Exercise Test (CPET)** |  |  |  |  |
| VO2max (L) | 1.86 (0.65) | 1.73 (0.56) | 1.89 (0.68) | 1.89 (0.65) |
| VO2max (ml/min/kg) | 24 (7) | 24 (6) | 25 (8) | 25 (7) |
| Workload (Watts) | 151 (50) | 142 (40) | 155 (53) | 153 (52) |
| O2 Pulse (ml/beat) | 10.6 (3.3) | 10.0 (2.8) | 10.8 (3.5) | 10.8 (3.4) |
| RER | 1.16 (0.08) | 1.16 (0.08) | 1.17 (0.07) | 1.15 (0.08) |
| **Strength Tests** |  |  |  |  |
| Isometric Knee Extension Torque (Nm) | 147 (57) | 137 (53) | 143 (54) | 156 (60) |
| Hand Grip Strength (kg) | 30 (11) | 29 (8) | 30 (11) | 31 (12) |
| <sup>1</sup> Mean (SD) |  |  |  |  |

No pre-exercise differences were found in participant characteristics across the three groups except HOMA-IR (Table 1). We observed relationships between some cardiometabolic health traits and circulating factors. As expected, adiposity measurements such as BMI and waist circumference (WC) were positively associated with higher blood pressure, whole-body and adipose-specific insulin resistance indices (HOMA-IR and Adipo-IR, respectively), and HbA1c (p<0.01; Figure S1A). WC was also positively associated with circulating branched-chain amino acids (BCAAs; isoleucine, leucine, and valine), which are known to be elevated in humans and rodents with insulin resistance and obesity^20^ (Figure S1A). More detailed clinical characteristics are available here (*REF: MoTrPAC Molecular Landscape, in review*).

To explore the relationship between baseline clinical characteristics and ASAT cell types, we performed a deconvolution analysis—estimating ASAT cell type proportions using a previously published ASAT single-nuclei (sn) RNAseq dataset and computational pipeline^21,22^ (Figure 1B, see ‘Deconvolution’ in Methods). As expected, the estimated proportion of macrophages was strongly correlated with higher adiposity, insulin resistance indices, and HbA1c, supporting the established link between ASAT macrophages and obesity and insulin resistance^23,24^ (p<0.001; Figure 1C). In contrast, the estimated proportions of adipocytes and vascular cells were negatively correlated with adiposity and insulin resistance indices, highlighting the complex role of ASAT cellular heterogeneity in cardiometabolic health traits (p<0.01; Figure 1C).

A significant degree of sexual dimorphism was observed across all ASAT omic layers measured at pre-exercise: 6150 transcripts (3039 up and 3111 down in males relative to females), 755 proteins (423 up and 332 down), 2133 phosphosites (1041 up and 1092 down), and 143 metabolites (36 up and 107 down) were significantly different between sexes (adjusted p<0.05; Figure 1D). To gain insight into sex-specific biological processes, we performed gene set enrichment analysis using CAMERA-PR^25^. At the transcriptomic level, females showed enrichment of lipid metabolism, branched-chain amino acid metabolism, and oxidative phosphorylation pathways, whereas males exhibited upregulation of protein translation and PDGFRβ signaling (adjusted p < 0.05; Figure 1E). Proteomic analysis revealed higher oxidative phosphorylation in females, while males displayed upregulation of pathways involved in actin cytoskeleton remodeling (e.g., actin filament, stress fiber, and myosin) and SUMOylation (Figure 1E). Kinase–substrate enrichment analysis (KSEA) of phosphoproteomic data identified sex-specific kinase activity, with all differentially enriched kinases upregulated in females. These included MAPK pathway kinases (e.g., PRKCH, RAF1, MAPK9, MAPK3, MAP3K8) and AKT signaling components (e.g., PDPK, GSK3A, GSK3B), both of which are known to strongly influence adipose cell proliferation, differentiation, and insulin signaling^26,27^ (Figure 1F). Metabolomic analysis showed higher abundances of phosphatidylethanolamines (PE) in females and increased saturated fatty acids (FA) in males (Figure 1G). Deconvolution analysis of the transcriptomics dataset suggested that the estimated proportion of Adipocyte-1, a subtype of mature adipocyte, previously characterized as ‘anti-oxidative’ with upregulation of genes related to anti-oxidative activity (*GPX1 & GPX4*)^21^, was higher in females compared to males (p=0.0007, Figure S1B). Conversely, the estimated proportion of vascular cells was lower in females (p=0.0003, Figure S1B), indicating intrinsic cellular heterogeneity by sex.

### Differentially expressed features following acute exercise

Differential analysis (DA) was performed across omic layers to investigate acute exercise effects. Importantly, we accounted for temporal changes observed in the non-exercising control (CON) group—where ASAT showed the most changes over time compared with muscle and blood even without exercise (*REF: MoTrPAC Molecular Landscape, in review*)—to better isolate the effects of acute endurance (EE) and resistance exercise (RE) (see ‘Differential Analysis’ in Methods). For example, in the CON group, 2801 transcripts (1306 upregulated, 1495 downregulated; Figure S2A) were differentially expressed at post 4hr, including core circadian clock genes *CLOCK*, *NR1D1/2,* and *PER1/2* (Figure S2B), despite the absence of any exercise stimulus. These genes have previously been reported as responsive to acute aerobic exercise in ASAT^10,12^ (Figure S2C). However, those earlier studies did not include non-exercising control groups, raising the possibility that the observed transcriptomic changes may have been at least partially driven by non-exercise factors, such as time of day, fasting, or biopsy-induced inflammation. Notably, temporal changes in these and other key clock genes like *CLOCK* were not differentially regulated across CON, EE, or RE (Figure S2B). Controlling for CON yielded markedly different outcomes across omic layers (Figure 2A), compared to differential analysis in one exercise group disregarding controls (Figure S2A). The number of differentially regulated transcripts peaked at post 45min (EE: n=57, RE: n=34), declined by post 4hr (EE: n=3, RE: 31), with these changes almost entirely returning to baseline by post 24hr (EE: n=3, RE: n=0), suggesting that transcriptional responses to acute exercise are largely transient (Figure 2A). Acute exercise triggered changes in proteins (EE: n=305, RE: n=2) and phosphosites (EE: n=280, RE: n=43) at post 4hr (adjusted p<0.05; Figure 2A). Differential metabolomic responses in ASAT also showed distinct temporal- and modality-specific patterns. Notably, the majority of differentially regulated metabolites were observed at post 4hr following EE (EE: n=93, RE: n=0) (adjusted p<0.05; Figure 2A). Principal component analysis (PCA) of log_2_fold changes further supported these distinct temporal and modality-dependent signatures in the transcriptome and metabolome (Figure 2B).

**Figure 2.**
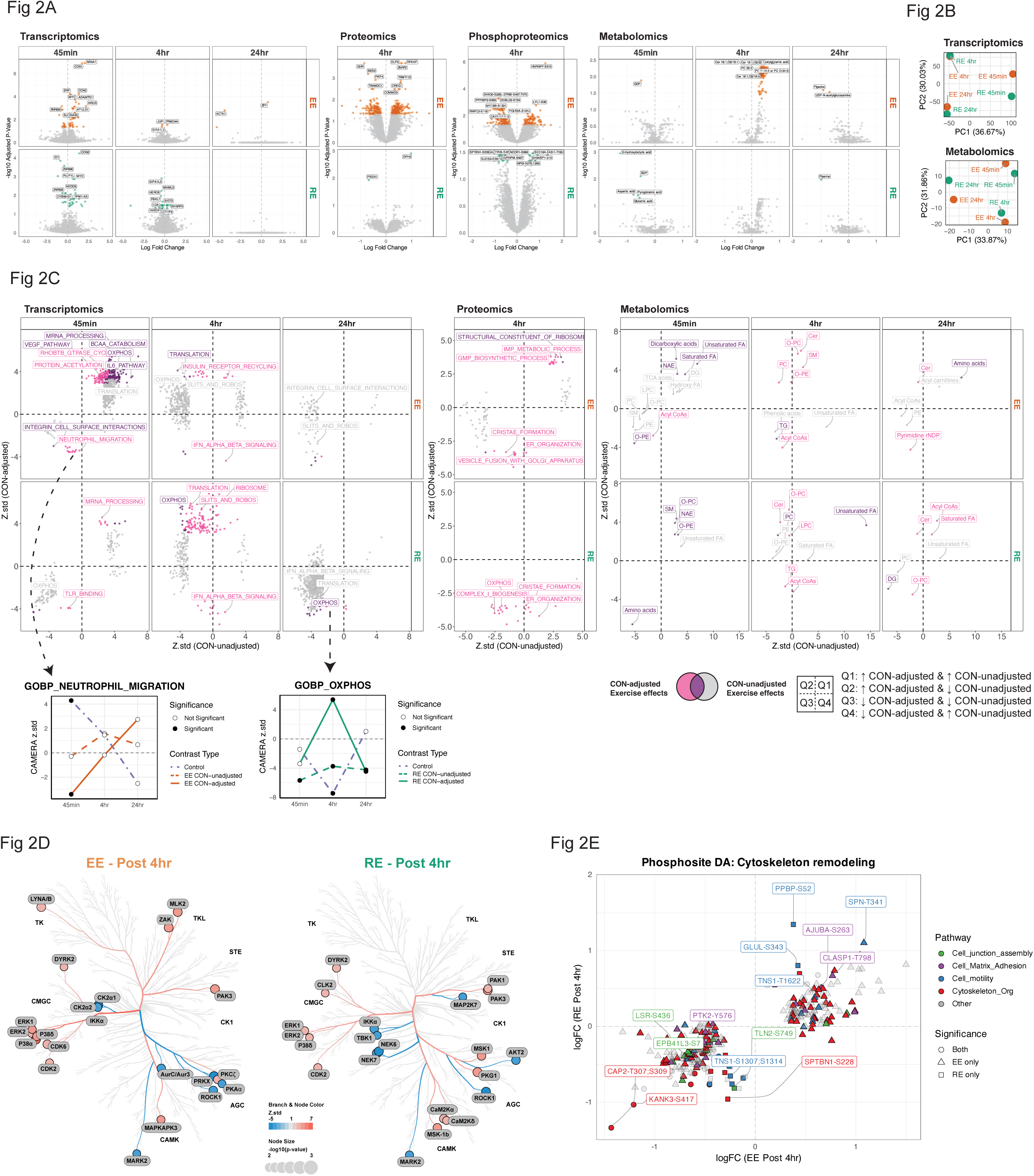
Multi-omic responses to acute exercise across modalities and time points. A) Volcano plots illustrating differentially regulated features across omic layers—transcriptomics, proteomics, phosphoproteomics, and metabolomics—stratified by exercise modality and time point. Differential analysis (DA) model accounts for changes observed in the control (CON) group. B) Principal component analysis of summarized log fold-change (logFC) values across modalities and time points, highlighting global shifts in transcriptomic and metabolomic responses to exercise. C) Scatter plots comparing significantly enriched biological terms from CAMERA-PR (adjusted p < 0.05) between CON-adjusted (y-axis) and CON-unadjusted (x-axis) DA results for transcriptomics, proteomics, and metabolomics. Hot pink indicates terms enriched only in CON-adjusted analyses; gray, terms enriched only in unadjusted analyses; purple, terms shared by both. Positive z.std values indicate upregulation; negative values indicate downregulation. D) Kinome phylogenetic tree annotated with enriched kinases (adjusted p < 0.1) from PTM-SEA at post 4Lhr following EE or RE. Red branches denote upregulated kinase activity; blue branches denote downregulation. See Methods for kinase group classification. E) Scatter plot of comparing logFC of differentially expressed phosphosites (adjusted p < 0.05) at post 4Lhr (EE or RE), annotated with cytoskeleton remodeling-related Gene Ontology terms. Shapes indicate whether a phosphosite was differentially regulated by EE only, RE only, or both.

Controlling for CON enabled more accurate pathway enrichment interpretation. Several pathways were upregulated at 45min post-EE in both adjusted and unadjusted models (e.g., ‘mRNA processing’, ‘VEGF signaling’, ‘BCAA catabolism’, and certain lipid classes like FA and dicarboxylic acids; Figure 2C, purple). However, CAMERA-PR applied to CON-adjusted DA revealed effects uniquely attributable to acute exercise when accounting for controls (Figure 2C, pink). For example, a reduction in neutrophil migration at 45 min post-EE was only evident after adjustment to the control group, which independently exhibited an increase in neutrophil migration thus masking the effect in the EE group without adjustment (Figure 2C). This distinction was even more pronounced at 4hr, where the directionality and magnitude of many enriched pathways were altered after CON adjustment. Notably, OXPHOS appeared downregulated at post 4hr in RE prior to CON adjustment, but this was primarily driven by stronger downregulation in the CON group; after adjustment, RE showed a relative upregulation of OXPHOS (Figure 2C). CON-adjusted enrichment of various lipid classes (e.g., TG, SM, CER, O-PC, O-PE) became evident post 4hr in both EE and RE (Figure 2C).

Focusing on CON-adjusted DA, upregulation of pathways related to angiogenesis, ECM remodeling, cell growth/differentiation, substrate utilization, and mitochondrial respiration at 45min post-EE, followed by protein translation at 4hr (adjusted p<0.05; Figure S2D). In RE, pathway enrichment predominantly emerged at 4hr and included mRNA processing, translation, ribosome biogenesis, substrate utilization, and mitochondrial respiration. By 24hr, most transcriptomic pathways were no longer significant, although some remained suppressed (translation in EE; mitochondrial biogenesis in RE). Proteomic analysis revealed suppression of substrate utilization and mitochondrial respiration pathways at 4hr in both EE and RE. In the metabolome, upregulation of FAs, N-acylethanolamines (NAEs), and dicarboxylic acids at 45min post-EE (adjusted p<0.05; Figure S2F). At 4hr, EE induced upregulation of sphingolipids (CER, SM) and glycerophospholipids (PC, LPC, O-PC, PE, O-PE), while suppressing acyl-CoAs and TGs. By 24hr, EE continued to alter lipid classes (LPC, PC, TG) and pyrimidine rNDPs (Figure S2F). RE upregulated glycerophospholipids at 45min, with some (e.g., O-PC, LPC) remaining elevated at 4hr and 24hr. Other lipid species (CER, LPC, DG, acyl-CoAs) were altered at 24hr post-RE, suggesting a prolonged metabolomic response.

PTM Signature Enrichment Analysis (PTMSEA) identified kinase activity differences at 4hr post-exercise (Figure 2D). Canonical MAPK pathway kinases ERK1/2 and P38, known to regulate adipogenesis and adipocyte structure^26^, were positively enriched in both EE and RE (adjusted p<0.1). Additional kinases enriched in EE or RE included those involved in cytoskeletal remodeling (e.g., PAK1/3, ROCK1, PKCζ, DYRK2, MARK2, CAMK2A/D, AURKC, LYN^28–31^). These kinase signatures occurred with enriched cytoskeletal pathways from overrepresentation analysis (ORA) of collapsed phosphoproteins (Figure S2G) and differentially expressed phosphosites mapping to processes such as ‘Cell junction assembly’, ‘Cell-matrix adhesion’, ‘Cell motility’, and ‘Cytoskeleton organization’ (Figure 2E). Several phosphosites in Tensin-1 (TNS1) were differentially phosphorylated or showed trends toward significance (Figure S2H). TNS1 is a scaffold protein anchoring actin filaments to integrins and mediates signal transduction between the ECM and cytoskeleton^32^. Notably, TNS1 has been reported as a downstream target of ROCK1^33^, and their protein abundances were significantly correlated in our data (p<0.001; Figure S2I), supporting the link between ROCK1 and TNS1 activation.

### Modality-specific multi-omic responses following acute exercise

Given the limited overlap in differentially expressed features between DA completed on EE and RE (Figure S3), we directly compared their effects on ASAT omic profiles to identify modality-specific responses (Figure 3A). Despite differences in exercise type, duration, and energy expenditure, the abundance of differentially expressed transcripts was minimal. Only one transcript, NR4A1 (also known as Nur77) — transcription activator that is known to regulate systemic glucose metabolism ^34^ and adipocyte differentiation ^35^— showed significantly greater upregulation in EE vs. RE at post 45min, with no significant differences at later time points (adjusted p<0.05; Figure 3B, C). The sphingomyelin species SM 42:2;O2 exhibited divergent responses at 45 minutes post-exercise, decreasing in EE but remaining unchanged in RE (Figure 3B, C). The reduced phosphorylation of NDRG1-S336 was more pronounced by RE at post 4hr. NDRG1 is required for the activation of a key regulator of adipogenesis, CEBPα, and its phosphorylation promotes adipocyte formation^36^. However, the functional role of the specific S336 phosphosite remains largely unexplored in the current literature. Meanwhile, only EE led to a robust increase in SLC44A1 (a choline transporter) protein abundance, which may explain the elevated PC species seen at post 4hr (Figure 2A).

**Figure 3.**
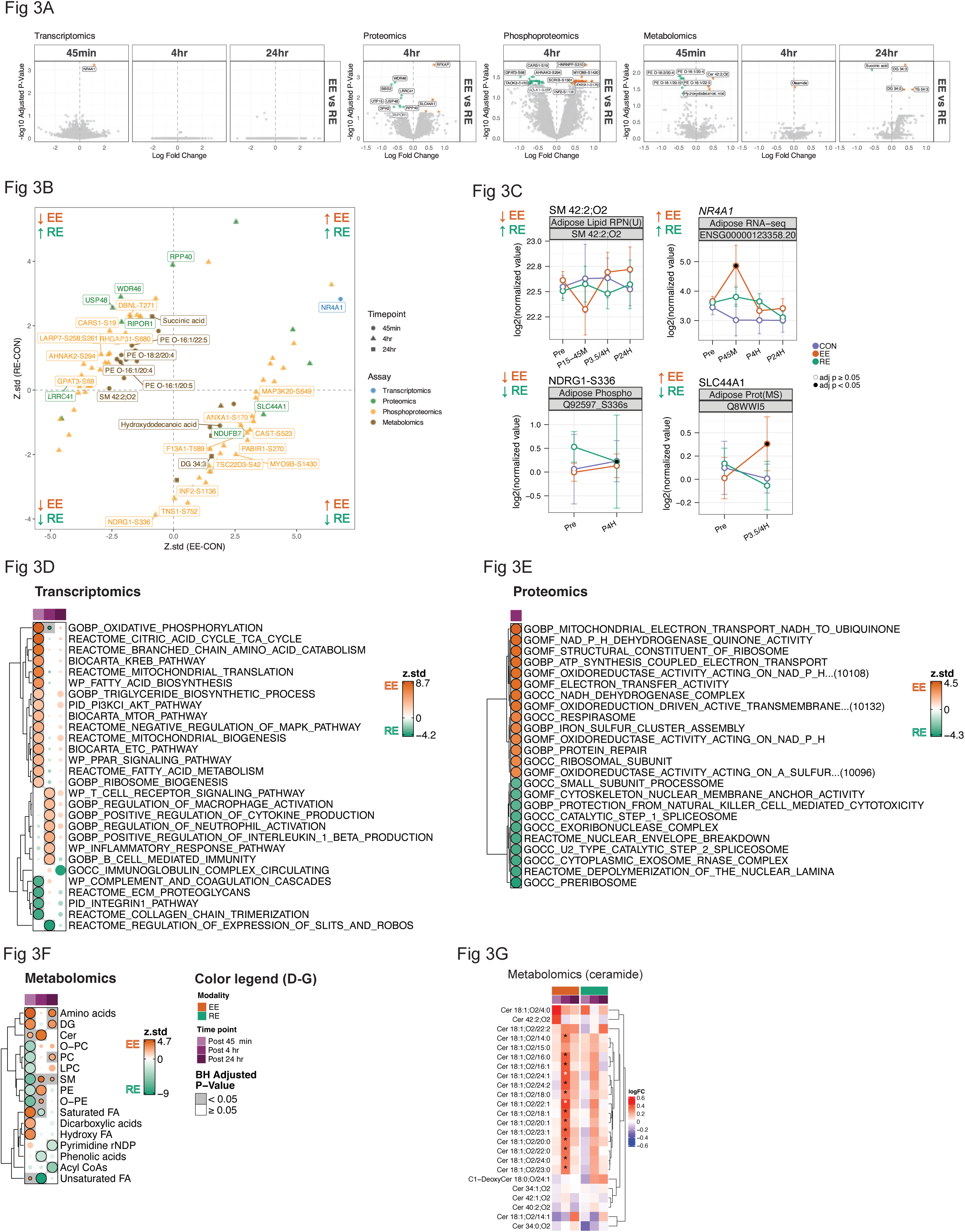
Acute exercise induces modality-specific responses in ASAT. A) Volcano plots illustrating differentially regulated features (adjusted p<0.05) between EE vs. RE across omic layers at each post-exercise time point. Positive logFC indicates greater enrichment in EE; negative logFC indicates greater enrichment in RE. B) Scatter plot comparing the z.std of differentially regulated features between EE and RE. Positive z.std values indicate upregulation; negative values indicate downregulation. Shape indicates time point. C) Individual line plots of features representing each quadrant of Figure 3B are shown. CAMERA-PR analysis comparing EE vs. RE in D) transcriptomic, E) proteomics, and F) metabolomics. Positive z.std values represent pathways more enriched in EE; negative values indicate enrichment in RE. G) Heatmap illustrating exercise responses in canonical ceramide species. *adjusted p<0.05.

Compared with RE, EE triggered greater early transcriptional enrichment of substrate metabolism, BCAA catabolism, ribosome biogenesis, oxidative phosphorylation, and MAPK signaling at post 45min, followed by immune cell activation and inflammatory responses at post 4 hr (adjusted p<0.05; Figure 3D). In contrast, RE induced greater enrichment in ECM signaling and remodeling pathways in post 45min, followed by Slits and Robos signaling (which has been partially linked to sympathetic activity and energetics in adipose^37^). Additionally, oxidative phosphorylation was induced at post 4hr with RE, and immunoglobulin complex at post 24 hr compared with EE (adjusted p<0.05; Figure 3D). The early transcriptional pathways enriched in EE vs. RE—particularly related to mitochondrial, ribosomal, and cytoskeletal structures (Figure 3D), were reflected in proteome-level enrichment at post 4hr (adjusted p<0.05, Figure 3E). RE, on the other hand, showed proteomic enrichment of the exosome complex, ribosome biogenesis, and spliceosome-related pathway at post 4 hr (adjusted p<0.05, Figure 3E). In metabolomics, EE induced greater alterations in amino acids, DG, ceramides, and fatty acids at post 45min, which then shifted toward changes in PE, SM, O-PE, and ceramide at post 4hr (adjusted p<0.05; Figure 3F). By post 24hr, amino acids, DG, and PC remained more significantly altered by EE vs. RE (adjusted p<0.05, Figure 3F). Conversely, RE induced greater changes in PC, O-PC, PE, LPC, SM, and O-PE at post 45 min compared with EE, followed by altered saturated FA and phenolic acids at post 4hr and pyrimidine rNDP at post 24 hr (adjusted p<0.05, Figure 3F). A notable EE-specific response was the increase in canonical ceramide species (C14:0–C24:0) at post 4hr (Figure 3G), aligning with upregulation of inflammatory pathways (Figure 3D). Acute inflammation is known to cause tissue ceramide accumulation through TNFa and TLR4 activation of ceramide synthesis^38^. However, ASAT ceramides did not remain elevated 24 hours after exercise, likely due to increased clearance, which would parallel results from acute exercise in skeletal muscle^39^. Enhanced clearance of ceramide post-exercise may perhaps result in lower ASAT ceramide content, which is associated with favorable cardiometabolic health^38^. Overall, these findings highlight the unique transcriptomic, proteomic, phosphoproteomic, and metabolomic responses induced by EE compared with RE, which may underlie modality-specific adaptations in ASAT by exercise training.

### Acute Exercise Induces Cell Type-Specific Transcriptional and Post-Translational Responses in ASAT

Given the heterogeneous relationship between ASAT cell types and cardiometabolic health traits observed in our deconvolution analysis (Figure 1C), we hypothesized that acute exercise would elicit distinct transcriptional responses across ASAT cell types. To test this, we curated a list of ASAT cell-type marker genes by integrating two of our previously published ASAT snRNAseq datasets^21,40^, capturing nine major cell types (see ‘Cell Type Analysis’ in Methods, see cell markers in Table S2). Cell type enrichment analysis of transcriptomic data using CAMERA-PR revealed modality- and time-dependent transcriptional activation across ASAT cell types (Figure 4A). Notably, at post 45 min, marker genes for Adipocyte-2—previously described as an insulin-responsive adipocyte subtype due to its upregulation of genes involved in lipolysis suppression (PDE3B), lipid metabolism (ABCA5), and insulin signaling^21^—were significantly upregulated in both training modalities (Figure 4A). In contrast, enrichment of Lipid-Associated Macrophages (LAM) was downregulated at post 45min in both modalities, suggesting early inhibition of pro-inflammatory macrophage activation following acute exercise. However, this suppression was reversed at post 4hr and post 24hr in response to EE, suggesting a delayed activation of macrophage-associated pathways (Figure 4A).

**Figure 4.**
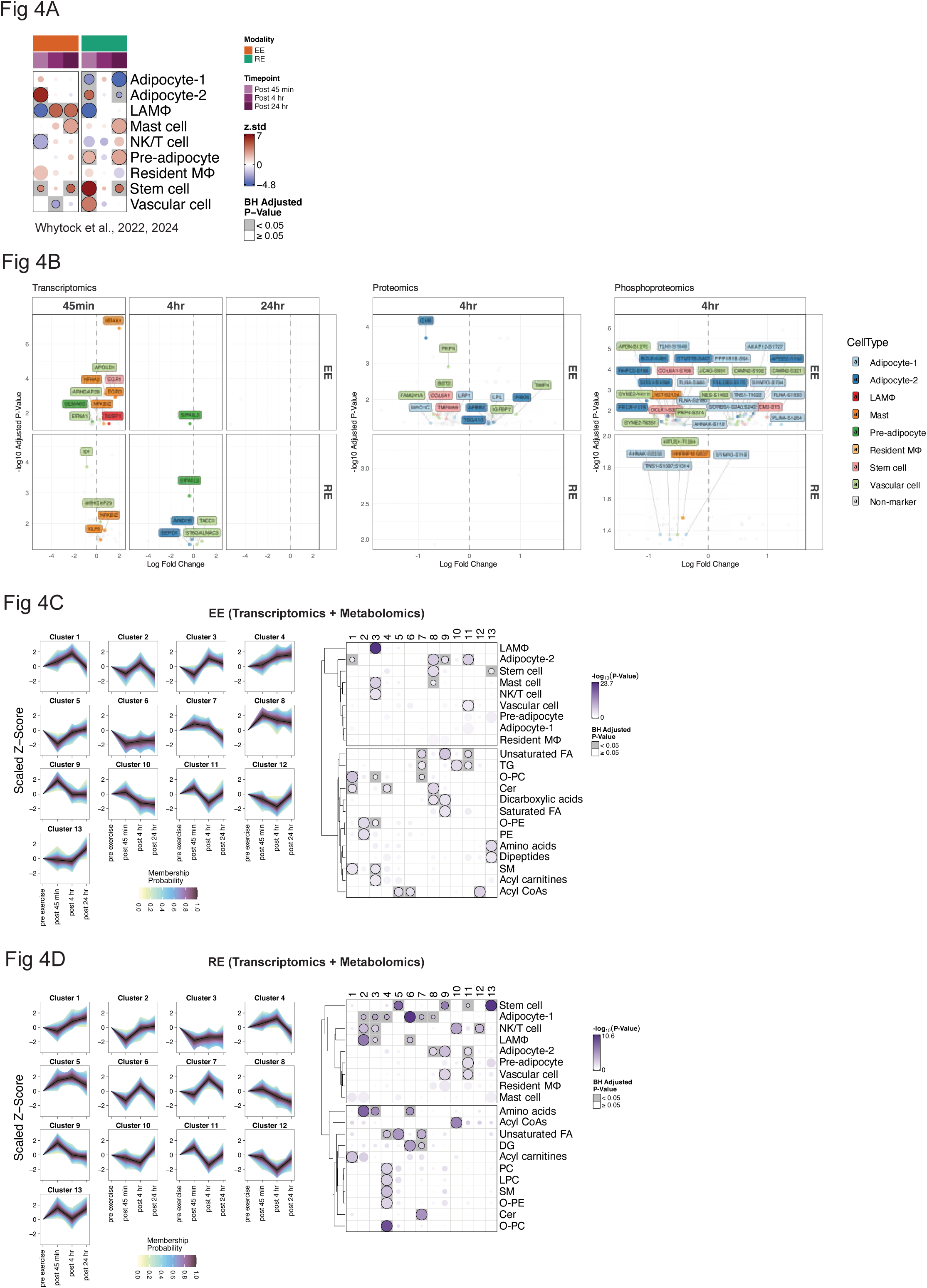
Acute exercise induces cell-type-specific responses in ASAT. A) Cell-type enrichment analysis using a reference cell signature matrix generated from ASAT single-nucleus RNA-seq datasets. CAMERA-PR was applied to ASAT transcriptomic data using the signature matrix as the gene set reference. B) Volcano plots of differentially regulated transcripts, proteins, and phosphosites mapped to known cell-type markers in EE and RE. ‘Non-marker’ indicates features not specific to any known ASAT cell type. C) EE-specific and D) RE-specific fuzzy c-means clustering analysis showing 13 trajectory clusters of combined object of transcriptomes and metabolomes. ORA was conducted on each cluster using the curated cell type markers and REFMET.

Additionally, transcriptional upregulated responses were observed in pre-adipocytes, stem cell, and vascular cell markers post 45min-RE, whereas enrichment of Adipocyte-1 was suppressed in response to RE, suggesting subtype-specific adipocyte responses occurring shortly after resistance exercise (Figure 4A). Furthermore, RE uniquely activated marker genes for both populations of mature adipocytes and progenitor cells (pre-adipocytes and stem cells) at post 24hr, indicating a long-lasting effect of RE on adipose progenitor and mature adipocyte transcriptional programs (Figure 4A). Importantly, despite these transcriptional changes, cell-type proportions assessed by deconvolution remained unchanged, suggesting that these transcriptional shifts may reflect activation states rather than changes in cell abundance (Figure S4A).

We next mapped differentially expressed features across omic layers to specific cell types. At 45 min, many transcripts were linked to mast cells and vascular cells (Figure 4B), even though these cell types were not enriched overall. In both proteomics and phosphoproteomics, differentially expressed features were primarily markers of adipocytes and vascular cells, suggesting these cells are major sites of functional change. Many differentially expressed phosphoproteins associated with cytoskeleton remodeling (e.g., SYNPO, FLNA, TNS1, TLN1) were attributed to Adipocyte-1 and vascular cells (e.g., SYNE2, NES, AFDN, PKP4), while metabolic regulators (e.g., PECR, BCL6) were linked to Adipocyte-2 (Figure 4B). Differentially phosphorylated sites between EE and RE were predominantly mapped to adipocytes at post 4 hr (Figure S4B). NR4A1 is known to activate mast cells, which release histamine^41^, a molecular transducer of exercise responses^42^ with wide-ranging beneficial effects in human skeletal muscle^43^. In ASAT, our data suggest that NR4A1 is a mast cell marker (Figure 4B, S4B). The upregulated gene expression of NR4A1 at post 45min-EE was followed by increased abundance of histamine in ASAT at post 4hr-EE (Figure S4C), suggesting the same exercise-induced histamine release — via mast cells — may therefore also be involved in responses of ASAT. To investigate whether exercise-induced changes in the ASAT metabolome mirror shifts in cell type activity, we performed fuzzy c-means clustering on a combined transcriptomics and metabolomics dataset. We identified 13 clusters per modality, each characterized by a distinct trajectory over time and unique patterns of cell type and metabolite class enrichment. For instance, in the EE group, Cluster 3 peaked at 4 hr post-exercise following a modest dip at 45 min, and was enriched for immune cell types including LAMs, mast cells, and NK/T cells, which aligns well with findings from the cell-type enrichment (Figure 4A). This immune-associated trajectory was accompanied by increased representation of metabolite classes such as O-PC, O-PE, and sphingomyelin, supporting the link between phospholipid metabolism and macrophages^44^ (Figure 4C). In RE, Clusters 2, 3, and 6 showed overlapping enrichment for LAMs and Adipocyte-1, with corresponding decreases post 45min—matching their transcriptional downregulation (Figure 4A). These clusters were also enriched for amino acids and diacylglycerols. Unsaturated fatty acids followed distinct temporal trajectories by modality: EE (Clusters 7, 9, 11) peaked at 45 min, while RE (Clusters 4, 5, 7) peaked at 4 hr. These clusters were associated with Adipocyte-2 in EE and Adipocyte-1 in RE (Figure 4C, D), suggesting modality-specific engagement of adipocyte subtypes in lipid metabolism. Similarly, ceramide-enriched clusters at post 4 hr exercise - cluster 1 in EE and cluster 7 in RE - were linked to Adipocyte-2 and Adipocyte-1, respectively, further indicating that EE and RE may differentially regulate lipid metabolism through distinct adipocyte subtypes.

#### Multi-Omic Network Analysis Links ASAT Features to Cardiometabolic Health and Exercise Responses

To explore the clinical relevance of multi-omic features in ASAT and their regulation by exercise, we applied weighted gene co-expression network analysis (WGCNA; Figure 5A)^45^ to identify co-regulated feature clusters (modules; Table S3) within each omic layer using baseline data: 13 transcriptomic modules (T0–T12), 15 proteomic modules (Pr0–Pr14), 19 phosphoproteomic modules (Ph0–Ph18), and 8 metabolomic modules (M0–M7) (Figure 5B; see WGCNA in Methods). ORA was performed to identify enriched pathways in each module. Next, we correlated module eigengenes—the first principal component summarizing overall module feature expression^46^—with cardiometabolic health traits and examined the module membership of features differentially regulated by exercise to assess the biological relevance of these modules. Several modules were significantly associated with adiposity (BMI, WC), insulin resistance (IR) indices (HOMA-IR, Adipo-IR), and HbA1c across omic layers, including T2, T5, Pr7, Pr12, Ph10, and M1 (Figure 5B), which showed high degrees of cross-omic intercorrelation (Figures S5A). Many of these modules were positively correlated with estimated macrophage proportion but inversely correlated with estimated adipocyte and vascular cell proportions (Figure S5B), and were enriched in pathways previously implicated in adipose-related metabolic complications, including inflammation (T2, Pr12)^47^, branched-chain amino acid (BCAA) metabolism (Pr7)^20^, post-transcriptional/translational modification (T5)^14^, cytoskeleton organization (Ph10)^48^, and ceramides and phosphatidylcholine (M1)^38^.

**Figure 5.**
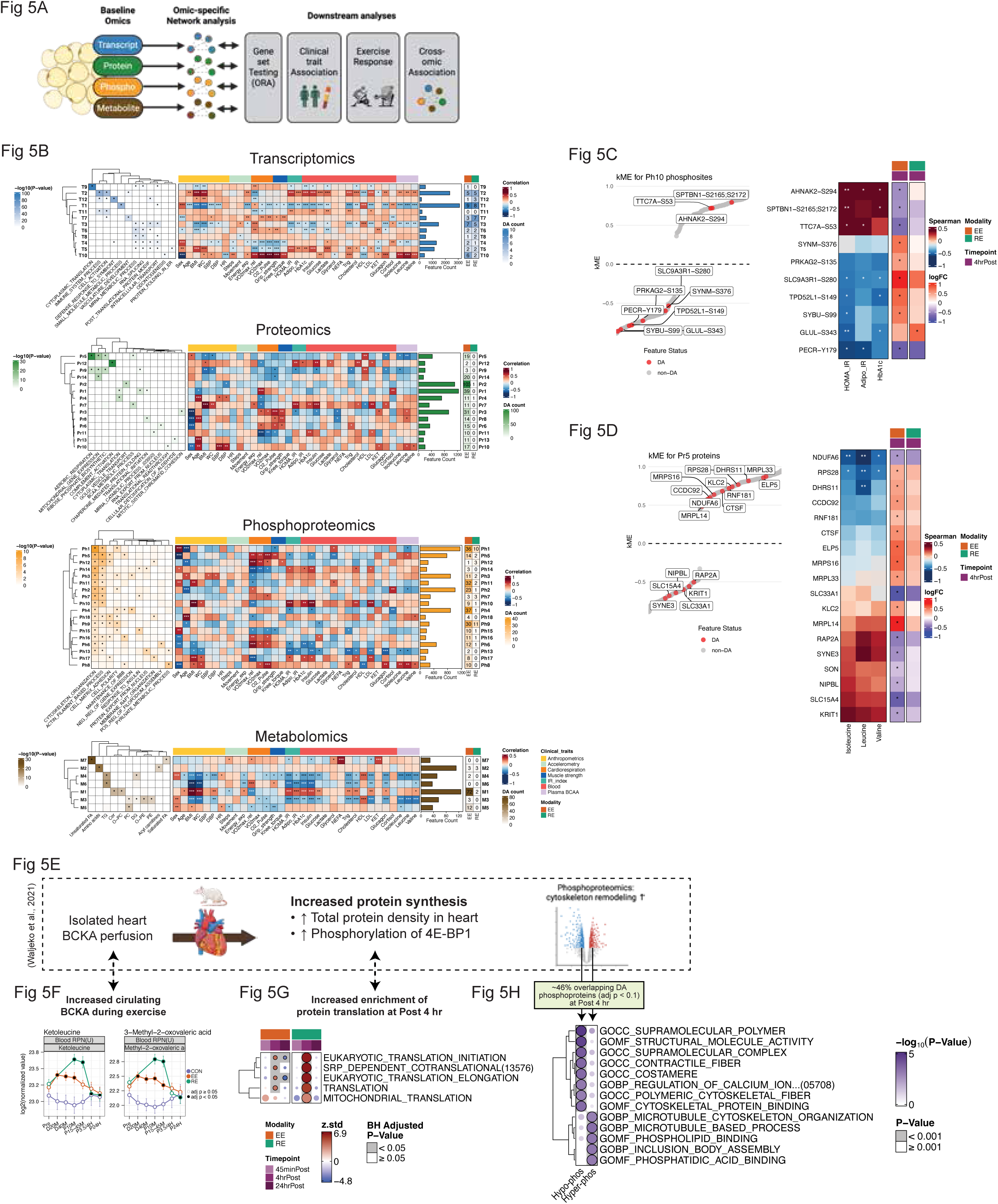
Acute exercise may induce metabolically favorable molecular responses in ASAT. A) Schematic overview of weighted gene co-expression network analysis (WGCNA) and downstream analyses. B) WGCNA was independently performed for each omic layer—transcriptomics, proteomics, phosphoproteomics, and metabolomics—using baseline data. Each panel includes: over-represented pathways per module (*FDR < 0.05), correlations between module eigengenes and z-scored clinical/subclinical traits (*p < 0.05; **p < 0.01; ***p < 0.001), module size (feature count), and the number of differentially regulated features (adjusted p<0.05) per module following acute exercise. C) Left: Scatterplot of module membership (kME) for phosphosites in Ph10; phosphosites differentially regulated by exercise are highlighted in red. Middle: Heatmaps showing baseline Spearman correlations between these phosphosites and clinical markers of insulin resistance (HOMA-IR, ADIPO-IR, HbA1c) (*p < 0.05; **p < 0.01). Right: Corresponding phosphosite fold changes after acute exercise (*adjusted p < 0.05). D) Left: Scatterplot of kME values for proteins in Pr5; red points indicate proteins differentially regulated by exercise. Middle: Heatmaps showing baseline correlations with circulating BCAAs (*p < 0.05; **p < 0.01). Right: Protein fold changes following exercise (*adjusted p < 0.05). E) Schematic summary of the BCKA perfusion experiment by Walejko et al., in which isolated rat hearts perfused with BCKAs via the Langendorff apparatus showed increased phosphorylation of 4E-BP1 (a translational repressor), enhanced protein synthesis, and activation of cytoskeleton-related phosphosignaling. F) Changes in plasma BCKA levels during and following EE and RE. G) CAMERA-PR enrichment results for ASAT transcriptomics filtered for protein translation-related pathways. H) ORA of differentially regulated phosphoproteins in ASAT at post 4Lhr, overlapping with phosphoproteins identified in the BCKA heart perfusion experiment. An adjusted p-value threshold of 0.1 was used for ASAT phosphosite selection. BCAA, Branched-chain amino acids; BCKA, branched-chain ketoacids.

We next examined whether acute exercise may impact features tightly associated with cardiometabolic health. Ph10 remained significantly associated with HOMA-IR and HbA1c after adjusting for key covariates (age, sex, WC, p<0.05, Figure S5C), suggesting a direct link between Ph10 and glycemic control. Among phosphosites differentially regulated by exercise in Ph10, those with positive intra-module connectivity (kME)—such as AHNAK2-S294 and SPTBN1-S2165;S2172—were positively correlated with IR indices and HbA1c, whereas phosphosites with negative kME—including SLC9A3R1-S280, TPH52L1-S148, and GLUL-S343—were inversely correlated with IR and HbA1c (p<0.05, Figure 5E, F). Strikingly, phosphosites with positive kME were downregulated, whereas those with negative kME were mostly upregulated, by acute exercise, particularly following EE at post 4 hr (adjusted p<0.05, Figure 5C). Exercise may therefore modulate phosphorylation of proteins involved in ASAT cytoskeletal organization acutely in a manner that contributes to favorable metabolic health.

BCAA catabolism is suppressed in ASAT in obesity, contributing to elevated circulating BCAA and IR^20,49^. In our data, module Pr5 was inversely correlated with circulating BCAAs (isoleucine, leucine, and valine) after adjusting for covariates (p < 0.05; Figure S5C). Within Pr5, proteins with negative kME values were all upregulated by EE at post 4 hr, whereas those with positive kME values were mostly downregulated (Figure 5D). Functional annotation of Pr5 highlighted enrichment for RNA processing, protein biosynthesis, cytoskeletal remodeling, and mitochondrial ribosomal components (Figure 5B), suggesting that acute exercise promotes proteomic remodeling in ASAT linked to these pathways.

In parallel, acute EE and RE significantly increased circulating levels of branched-chain keto acids (BCKAs)—specifically ketoleucine and 2-keto-3-methylvalerate—which are derived from BCAAs through transamination by branched-chain amino transaminase (Figure 5F). Notably, prior work by Walejko et al. showed that acute exposure of perfused, isolated working hearts to elevated BCKA levels (mirroring those seen in obesity) was sufficient to induce phosphoprotein changes, cytoskeletal remodeling, and activation of protein synthesis^50^ (Figure 5E). Strikingly, our results echo these findings: translational pathways were upregulated at post 4 hr in both EE and RE (Figure 5G), and nearly half of the phosphoproteins differentially phosphorylated in the BCKA-exposed heart overlapped with those regulated by exercise in ASAT. These overlapping phosphoproteins were enriched for cytoskeleton remodeling functions (Figure 5H), consistent with the remodeling signatures reported by Walejko et al. Together, this supports the hypothesis that transient, exercise-induced increases in BCKAs may trigger shared phosphoproteomic and cell structural remodeling programs in adipose tissue. While chronic elevation of BCKAs—as seen in obesity—may contribute to maladaptive cardiac hypertrophy through sustained activation of protein synthesis, acute exercise-induced BCKA elevations may be health-promoting by virtue of their intermittent rather than chronic effects.

### Adipose Tissue as a Source of Exercise-Induced Exerkines

Acute exercise can stimulate the release of exerkines—secreted factors that mediate metabolic and functional adaptations in distant tissues—with ASAT serving as a key contributor^19^. To characterize the exercise-responsive secretome of ASAT, we filtered differentially expressed transcripts and proteins using curated secretome resources, including MoTrPAC’s blood proteomics panel (*REF: MoTrPAC Blood Companion, in review*) and The Human Protein Atlas^51^, which includes 3,277 known secreted proteins. This yielded 60 unique candidate proteins (Figure 6B, S6A), the majority of which were identified through proteomics and induced at 4 hr post-exercise (Figure S6A, Table S4). For metabolites, 92 differentially expressed ASAT features overlapped with plasma metabolomics (Figure 6C). At baseline, ASAT abundances of TIMP4 and CCN1 were significantly correlated with their circulating protein levels (p < 0.05; Figure S6B), suggesting that ASAT may contribute to their systemic concentrations at rest. Likewise, ASAT-derived glycerophospholipids (e.g., PC, PE) and sphingolipids (e.g., SM, ceramide) showed strong correlations with their plasma counterparts (p < 0.05; Figure S6C).

**Figure 6.**
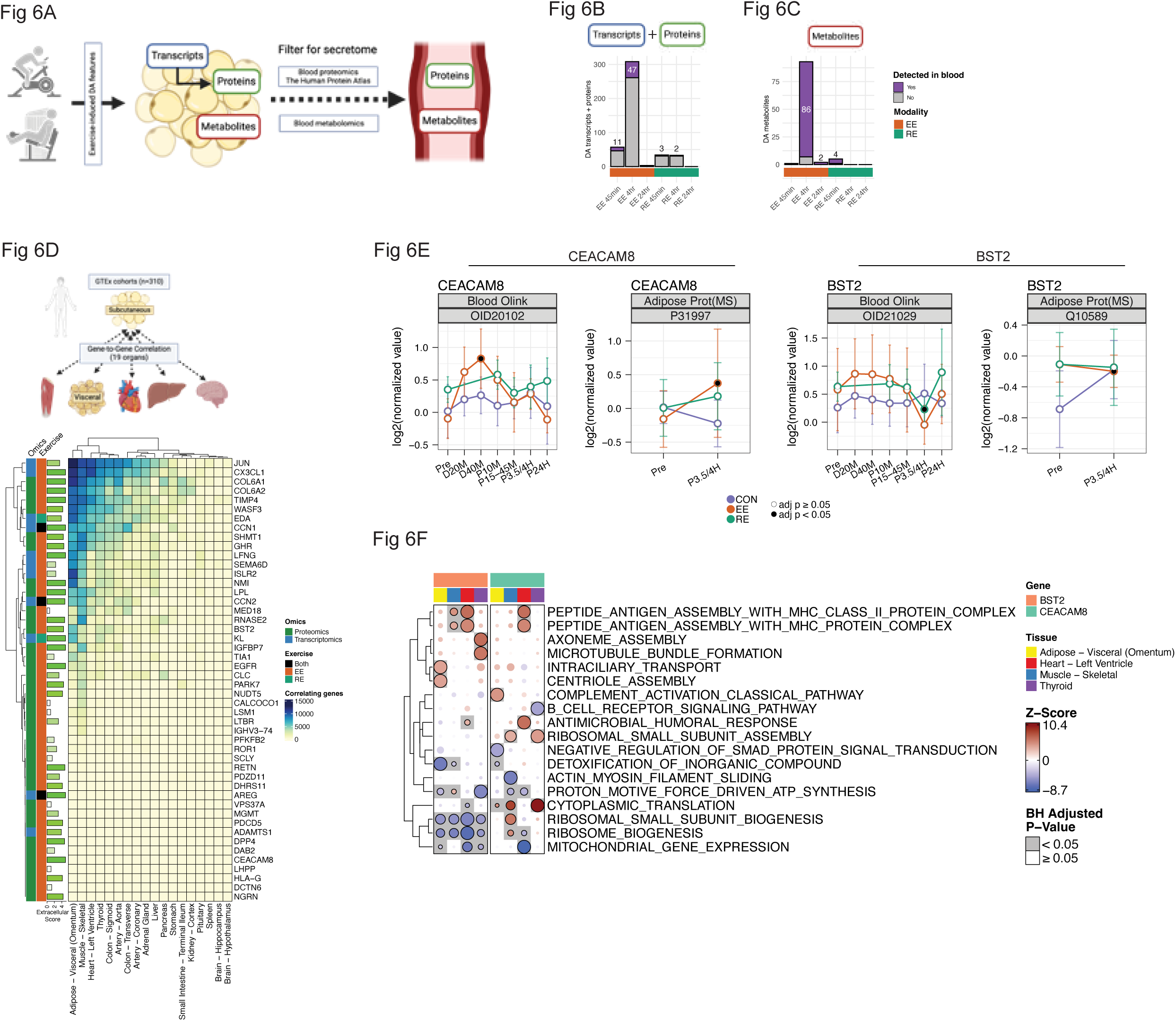
Identification of ASAT-secreted exerkine candidates and their potential endocrine role. A) Schematic overview of the secretome screening pipeline. Transcripts and proteins differentially regulated at any post-exercise timepoint (post 45 min, 4r, and 24hr) were filtered using MoTrPAC blood Olink proteomics data and secretome annotations from the Human Protein Atlas. Metabolites were filtered against MoTrPAC blood metabolomics. B) Bar plots showing the number of differentially regulated transcripts and proteins per time point. Purple bars indicate features annotated as secreted. C) Bar plots showing the number of differentially regulated metabolites per time point. Purple bars indicate metabolites also detected in plasma. C) Integration with GTEx: RNA-seq from n = 310 GTEx subjects across 18 solid tissues was used to correlate 48 GTEx-mappable ASAT-exerkine candidate genes with genes in distal tissues. The heatmap displays the number of significantly correlated genes (adjusted p<0.05) per exerkine candidate per target tissue. Green bars indicate the extracellular region score (range 0–5) from COMPARTMENTS, reflecting the strongest extracellular annotation across multiple categories (region, space, exosome, vesicle). E) Exercise responses of CEACAM8 and BST2 proteins in both ASAT and circulation. F) GD-CAT analysis showing pathways associated with SAT-CEACAM8 and SAT-BST2 in selected target tissues. CAMERA-PR was used for enrichment testing.

To explore potential endocrine roles of ASAT-derived exerkines, we applied inter-tissue gene-gene correlation analysis using GTEx—the most comprehensive multi-tissue human transcriptomics resource^52^ (see Gene-to-Gene Correlations in Methods). We also assigned each transcript an “extracellular score (0-5)” using COMPARTMENTS (see ‘Extracellular score in Methods’)^53^ to estimate its secretory likelihood. Secreted candidate genes expressed in SAT displayed distinct correlations with transcriptional programs in distant tissues—most notably visceral adipose tissue, skeletal muscle, heart, and thyroid (adjusted p<0.05, Figure 6D). To infer potential functional pathways in recipient tissues, we applied Gene-Derived Correlations Across Tissues (GD-CAT)^54^, which assumes that if a gene encoding a secreted protein in one tissue correlates with specific transcriptional programs in another, the protein may mediate inter-tissue crosstalk^55^. We focused on two exerkine candidates significantly altered in plasma during or after exercise: *CEACAM8* (CD66b) and *BST2* (adjusted p<0.05, Figure 6E). *CEACAM8*, a marker gene for LAM (Figure 4C) and known for its immunoregulatory role^56^, had a high extracellular score of 5 and was acutely increased in the plasma during EE, with increased protein abundance in ASAT at post 4hr-EE (Figure 6E). In GTEx, SAT-*CEACAM8* was associated with upregulated immune responses (e.g., MHC assembly, antimicrobial humoral response) and downregulated ribosome biogenesis in visceral adipose tissue (VAT) and thyroid (adjusted p<0.05, Figure 6F, Table S5). In contrast, SAT-CEACAM8 correlated with enhanced protein translation and dampened B cell receptor signaling in skeletal muscle, suggesting possible tissue-specific endocrine functions (adjusted p<0.05, Figure 6F). BST2, a transmembrane glycoprotein linked to obesity-associated inflammation^57^ with extracellular score of 4.5, decreased systemically at post 4hr-RE (adjusted p<0.05). Interestingly, this coincided with the prevention of an increase in ASAT BST2 post-exercise (EE, adjusted p<0.05; RE, adjusted p=0.14), contrasting with its rise in CON (Figure 6E). In GTEx, SAT-BST2 was linked to downregulated mitochondrial energetics (proton motive force-driven ATP synthesis) in VAT, heart, and thyroid, but not skeletal muscle (adjusted p<0.05, Figure 6F, Table S5). Conversely, SAT-BST2 was associated with upregulated complement activation in skeletal muscle (adjusted p<0.05, Figure 6F). These data suggest that exercise-induced reductions in BST2 may confer systemic metabolic benefits by limiting inflammation and preserving mitochondrial function across tissues. Together, these findings reveal novel ASAT-derived exerkine candidates modulated by acute exercise and demonstrate how integrated transcriptomic correlations can uncover potential endocrine effects of adipose tissue on distant organs.

### Exercise-Induced ASAT Secretomes: Adaptations in the Endocrine Network with Exercise Training

To investigate whether ASAT-derived exerkine candidates contribute to systemic adaptations during endurance training, we leveraged the MoTrPAC rat training dataset^58^, which includes transcriptomic profiles of subcutaneous white adipose tissue (WATSC) and 17 other tissues at multiple time points (1, 2, 4, and 8 weeks; TR1W–TR8W), along with age-matched non-trained controls (CON). This dataset enabled us to assess how WATSC exerkine candidates correlate with target tissues and how these relationships evolve with training. We applied Quantitative Endocrine Network Interaction Estimation^59^, a systems genetics framework that computes a “secretome score” (Ssec) to estimate the endocrine influence of secretory genes across origin–target tissue pairs. Briefly, for each gene in the origin tissue, a -log10 sum of the correlation p-values with all genes in the target tissue is computed to generate Ssec, where higher scores indicate broader or stronger endocrine connectivity (see QENIE in Methods). Using this framework, we previously identified WATSC as one of the most responsive endocrine tissues to endurance training^60^. We designated WATSC as the origin tissue and calculated Ssec for 2,248 secretory protein-coding genes found in humans (Figure 6A) across multiple target tissues (Figure 7A; see Methods; full dataset in Table S6). Given the strong sexual dimorphism of WATSC in this cohort^61^, all correlations were adjusted for sex. Ssec distributions revealed tissue- and time-dependent endocrine connectivity changes (Figure 7B). Out of 60 ASAT protein exerkine candidates we identified (Figure 6), 53 of them were mappable in rat WATSC. Focusing on these 53 candidates, we found significant alteration in their Ssec rank distributions between control and 8-week trained rats for key endocrine connections: WATSC-to-vastus lateralis, liver, and kidney (adjusted p<0.05, Figure 7C). These rank shifts were paralleled by increases in average Ssec values in 8-week trained rats compared to controls in all three tissue connections (adjusted p<0.05, Figure S7A), indicating strengthened endocrine influence of WATSC with training.

**Figure 7.**
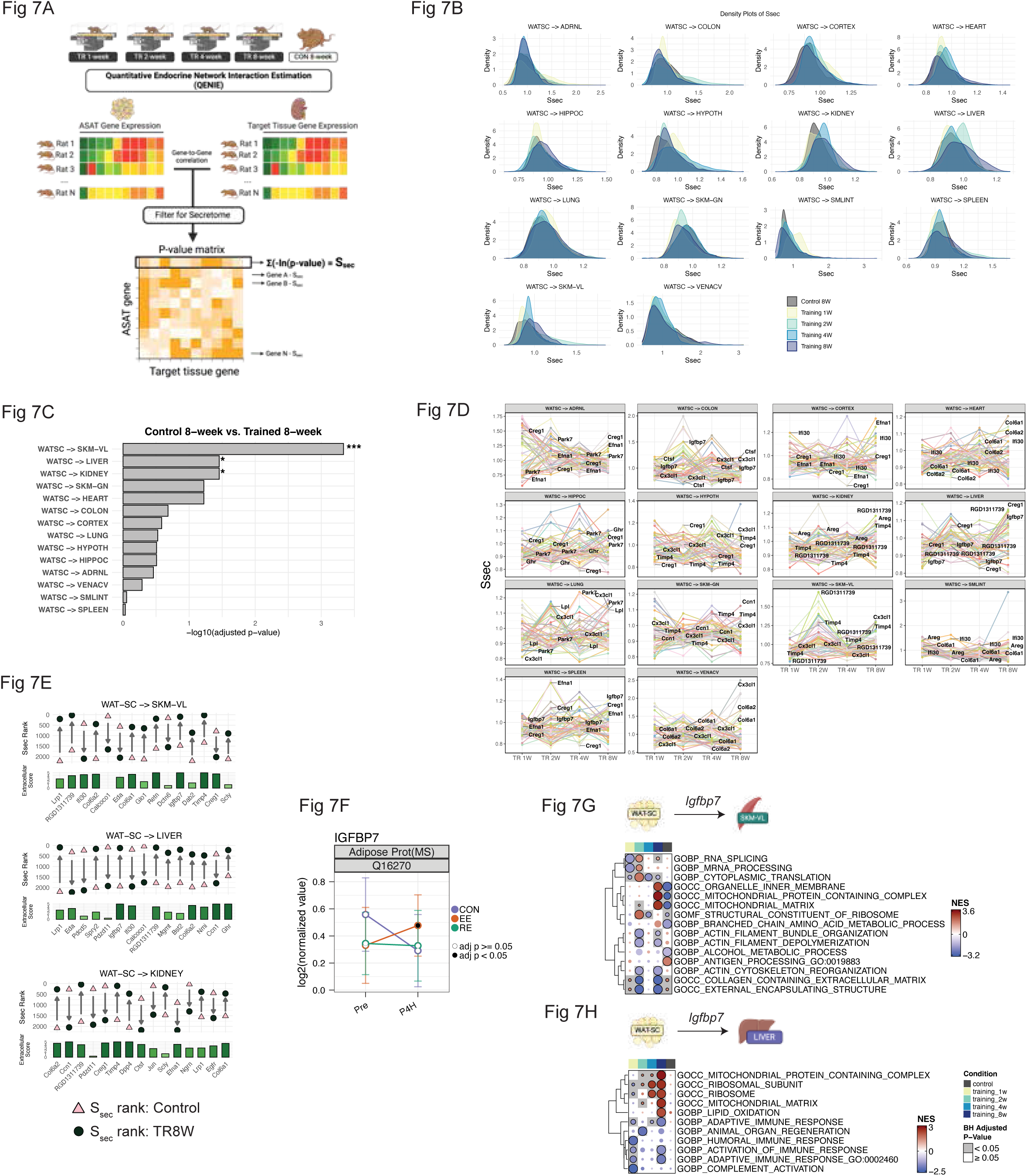
Contribution of ASAT-exerkine candidates to training-induced remodeling of adipose tissue endocrine network. A) Schematic workflow for applying QENIE to the MoTrPAC rat endurance training study. Human exerkine candidates were mapped to rat orthologs, and the same secretome filters as in Figure 6 were applied, yielding 2,248 WATSC secretory genes. B) Density plots showing distributions of secretome scores (Ssec) across WATSC-to-target tissue connections over 1, 2, 4, and 8 weeks of endurance training (TR1W, TR2W, TR4W, TR8W), compared with non-trained control rats (CON). C) Wilcoxon signed-rank test comparing Ssec rank distributions of 53 mappable ASAT-exerkine candidates between CON and TR8W for each WATSC-to-target tissue connection. Adjusted **p < 0.01, ***p < 0.001, **p < 0.0001. D) Time-course trajectories of Ssec values for the 53 exerkine candidates across training weeks. Top 3 genes per tissue connection at TR8W (with extracellular score ≥L4) are labeled. E) String plots showing the top 15 exerkine candidates with the greatest Ssec rank shift between CON and TR8W for WATSC-to-vastus lateralis, liver, and kidney connections. Pink triangle, CON; dark green circle, TR8W. Vertical green bars denote extracellular scores. F) Exercise-induced response of IGFBP7 protein at post 4Lhr. G) FGSEA results showing pathways associated with WATSC-Igfbp7 in vastus lateralis. H) FGSEA results showing pathways associated with WATSC-Igfbp7 in the liver. ADREN, adrenal gland; HIPPO, hippocampus; HYPOTH, hypothalamus; SKM-GN/VL, skeletal muscle gastrocnemius/vastus lateralis; SMLINT, small intestine; VENACV, vena cava; NES, normalized enrichment score.

Time-course analysis of individual Ssec trajectories showed progressive increases for several exerkines. For example, *CX3CL1* (fractalkine)—a known exerkine also released from skeletal muscle^62^—displayed increased Ssec over time, peaking at 8 weeks in WATSC-to-colon, hypothalamus, and lung connections (Figure 7D). Feature-level comparisons revealed heterogeneous adaptations across candidates and tissues (Figure S7B). While overall Ssec ranks for WATSC-to-vena cava were unchanged (Figure 7C), several candidates (e.g., *TIMP4, LHPP, PFKFB2, CX3CL1*) showed dramatically elevated Ssec values at 8 weeks of training, suggesting a role in vascular crosstalk during habitual exercise adaptation (Figure S7B). We next highlighted top candidates showing the greatest increases in Ssec rank between control and 8-week trained rats across the vastus lateralis, liver, and kidney (Figure 7E). *IGFBP7* (Insulin-like Growth Factor Binding Protein 7), which was induced by EE at 4 hr (Figure 7F), emerged as a notable candidate in WATSC-to-vastus lateralis and WATSC-to-liver connections (Figure 7E). In the WATSC-to-vastus lateralis axis, *IGFBP7* rose from rank 1477 in CON to 95 in trained rats. Similarly, in the WATSC-to-liver axis, *IGFBP7* jumped from 1942 to 121 (Figure 7E). GD-CAT analysis revealed that *IGFBP7* was associated with enhanced mitochondrial metabolism pathways in the vastus lateralis of 8-week trained rats, while control rats showed enrichment of inflammatory signaling and mitochondrial suppression (adjusted p<0.05; Figure 7G). These findings align with *in vitro* evidence showing improved mitochondrial respiration in myotubes treated with IGFBP7-conditioned media^63^, supporting its potential endocrine role in exercise adaptations. In the liver, WATSC-*IGFBP7* correlated with upregulated mitochondrial complex expression, translation, and lipid oxidation, along with downregulated immune responses in 8-week trained rats (adjusted p<0.05; Figure 7H)—associations absent in control rats. Notably, IGFBP7 is a known tumor suppressor; its deletion or downregulation in the liver induces hepatocellular carcinoma^64,65^, raising the possibility of additional protective roles in exercise-adapted crosstalk mediated by WATSC.

Together, these systems genetics analyses suggest that acutely-induced ASAT exerkines can exert tissue-specific endocrine effects, and that repeated endurance training progressively reshapes the WATSC-mediated secretome landscape. This endocrine network remodeling may underlie key systemic adaptations induced by habitual exercise.

## DISCUSSION

In this study, we captured modality- and time-specific multi-omic molecular responses in human abdominal subcutaneous adipose tissue (ASAT) following acute exercise using both endurance (EE) and resistance (RE) modalities across multiple post-exercise time points in sedentary adults. Acute exercise triggered diverse transcriptional and proteomic responses involving substrate metabolism, angiogenesis, adipogenesis, ECM remodeling, protein synthesis, cell signaling, cytoskeletal remodeling, and inflammation and metabolite-specific changes. Using integrative bioinformatics, we identified cell-type-dependent responses, novel exercise-responsive features linked to cardiometabolic health, and a set of putative adipose-derived exerkines. In addition to these post-acute-exercise changes, baseline sex differences and variations in cell-type proportions were examined, offering insights into how acute exercise responses may initiate early adaptations towards long-term benefits.

Adipose tissue exhibits circadian rhythmicity in gene expression patterns and physiological functions. This includes mediating key metabolic enzymes and transport systems involved in lipid metabolism, lipolysis/lipogenesis and the secretion of adipokines, thus influencing systemic metabolism, energy homeostasis and inflammatory processes^66,67^. By inclusion of a non-exercising control group in our differential analysis, we were able to dissociate exercise-specific effects from confounding time-of-day and fasting influences—particularly pronounced in adipose tissue and more substantial than anticipated (*REF: MoTrPAC Molecular Landscape, in review*). Although previous studies have reported post-acute-exercise changes in core clock genes^10,68^, we observed similar patterns in non-exercising controls, suggesting that some of these transcriptomic shifts may be driven by time-of-day effects rather than exercise *per se*. This comparison provides an additional layer of confidence in interpreting true exercise-responsive signals. Our design also enabled us to detect exerkines such as BST2, whose significance emerged specifically in relation to the control group trajectory. To our knowledge, this study represents one of the first efforts to directly account for non-exercise-mediated changes when assessing acute exercise-induced molecular responses in adipose tissue and may serve as a reference point for future studies aiming to disentangle these overlapping influences.

The actin cytoskeleton plays a central role in insulin action and adipocyte size dynamics^48,69^. Although the precise functions of many phosphosites remain unknown, several within the Ph10 module were strongly associated with insulin resistance. Notably, acute exercise modulated these phosphosites in a direction consistent with improved metabolic health—sites positively associated with insulin resistance tended to be downregulated, while those inversely associated with insulin resistance were upregulated. The overall pattern suggests that acute exercise may activate phosphosignaling within cytoskeletal remodeling pathways that contribute to favorable metabolic responses. Cell-type analysis further revealed that Adipocyte-1—characterized by anti-oxidative and translational activity—and vascular cells were the predominant sources of cytoskeleton-related phosphosignaling. This raises the possibility that these cell types may be key mediators of early, metabolically beneficial responses to acute exercise.

A major advance of this study was the identification of adipose-derived exerkine candidates, derived by filtering differentially regulated features through plasma proteomics, metabolomics, and curated secretome databases. Using GD-CAT^54^ and GTEx pan-tissue transcriptomics^52^, we inferred potential endocrine actions of several candidates—such as CEACAM8 and BST2—whose predicted effects varied across tissues. Further, by applying QENIE^59^ to the MoTrPAC rat endurance training dataset^58^, we found that adipose endocrine networks are remodeled with training. Several exerkine candidates, particularly IGFBP7, appeared central to this adaptive rewiring, especially in WATSC-to-vastus lateralis and WATSC-to-liver axes. Interestingly, 8 weeks of endurance training led to modest but significant downregulation of *IGFBP7* transcript levels in WATSC (male rats: –0.11 log_₂_FC; female rats: –0.22 log_₂_FC; FDR = 0.008)^58^, without corresponding changes in protein abundance—suggesting that, like many exerkines^70,71^, the systemic effects of IGFBP7 may be primarily driven by acute induction rather than chronic upregulation. While our systems genetics approach is correlation-based and requires experimental validation, this framework has been successfully used to uncover novel crosstalk regulators that have been validated in pre-clinical models^55,59,72^. Additionally, other candidates—TIMP4 (Tissue Inhibitor of Metalloproteinase 4), LRP1 (Low-density lipoprotein receptor-related protein 1), and RETN (Resistin)— also emerged among top-ranking factors across several adipose-to-target tissue connections in trained rats, warranting further mechanistic investigation.

Despite the strengths of our design and depth of analysis, several limitations should be acknowledged: Participants served as their own control within each exercise group, but comparisons across groups were not conducted using a cross over design. By design the two exercise modalities were not matched for exercise duration and intensity or energy expenditure^73^. While this limits certain interpretations, the goal was to model real-world clinical approaches rather than creating tightly matched interventions. Moreover, proteomic and phosphoproteomic profiling was not performed at post 45 min or post 24 hr, possibly resulting in missed windows of dynamic regulation. Given the robust changes at post 4hr, additional time points could provide a more complete temporal map. Finally, our secretome and endocrine modeling leveraged rodent training data, which may not fully translate to humans. Integration of anticipated MoTrPAC 12-week training data in sedentary adults will allow deeper analysis of secretome dynamics and tissue crosstalk in the context of long-term adaptation.

## ACKNOWLEDGEMENTS

We would like to acknowledge Dr. Marcus Seldin at the University of California, Irvine, for assisting GTEx and QENIE analysis. The graphical abstract and schematic figures were created using BioRender (www.biorender.com), and confirmation of publication and licensing rights was obtained. The MoTrPAC Study is supported by NIH grants U24OD026629 (Bioinformatics Center), U24DK112349, U24DK112342, U24DK112340, U24DK112341, U24DK112326, U24DK112331, U24DK112348 (Chemical Analysis Sites), U01AR071133, U01AR071130, U01AR071124, U01AR071128, U01AR071150, U01AR071160, U01AR071158 (Clinical Centers), U24AR071113 (Consortium Coordinating Center), U01AG055133, U01AG055137, U01AG055135, U01AG070959, U01AG070960, and U01AG070928 (Pre-Clinical Animal Sites).

Additional grant funding: D.H.K: K23HL164980, 23CDA1040581; N.M and S.E: P30AG094848.

## AUTHOR CONTRIBUTIONS

JNA, MJW, and LMS conceptualized and designed the study. CA, CAJ, and TJS performed data analyses and developed figures. CA, CAJ, KLW, GMM, JAS, ZH, MRV, JN, SEE, NM, YS, MFP, SCB, LJG, KAE, CBN, BCB, JNA, MJW, and LMS contributed to data interpretation. CA and LMS drafted the work with input from other authors. All authors have participated in revising the work. All authors have read and approved the final version of the manuscript.

## MoTrPAC Study Group

**Bioinformatics Center**: David Amar, Trevor Hastie, David Jimenez-Morales, Daniel H. Katz, Malene E. Lindholm, Samuel Montalvo, Robert Tibshirani, Jay Yu, Jimmy Zhen, Euan A. Ashley, Matthew T. Wheeler

**Biospecimens Repository:** Sandra T. May, Jessica L. Rooney, Russell Tracy

**Data Management, Analysis, and Quality Control Center:** Catherine Gervais, Fang-Chi Hsu, Byron C. Jaeger, David Popoli, Joseph Rigdon, Courtney G. Simmons, Cynthia L. Stowe, Michael E. Miller

**Exercise Intervention Core:** W. Jack Rejeski

**NIH:** Ashley Y. Xia

**Preclinical Animal Study Sites:** Sue C. Bodine, R. Scott Rector

**Chemical Analysis Sites:** Hiba Abou Assi, Mary Anne S. Amper, Brian J. Andonian, Isaac K. Attah, Jacob L. Barber, Kevin Bonanno, Clarisa Chavez Martinez, Natalie M. Clark, Johanna Y. Fleischman, David A. Gaul, Yongchao Ge, Marina A. Gritsenko, Joshua R. Hansen, Patrick Hart, Zhenxin Hou, Chelsea M. Hutchinson-Bunch, Olga Ilkayeva, Gayatri Iyer, Pierre M. Jean-Beltran, Christopher A. Jin, Maureen T. Kachman, Hasmik Keshishian, Damon T. Leach, Minghui Lu, D. R. Mani, Gina M. Many, Nada Marjanovic, Nikhil Milind, Matthew E. Monroe, Ronald J. Moore, Venugopalan D. Nair, German Nudelman, Nora-Lovette Okwara, Vladislav A. Petyuk, Paul D. Piehowski, Hanna Pincas, Wei-Jun Qian, Prashant Rao, Abraham Raskind, Alexander Raskind, Stas Rirak, Jeremy M. Robbins, Margaret Robinson, Tyler J. Sagendorf, James A. Sanford, Gregory R. Smith, Kevin S. Smith, Yifei Sun, Mital Vasoya, Nikolai G. Vetr, Alexandria Vornholt, Yilin Xie, Xuechen Yu, Elena Zaslavsky, Zidong Zhang, Bingqing Zhao, Joshua N. Adkins, Charles F. Burant, Steven A. Carr, Clary B. Clish, Facundo M. Fernandez, Robert E. Gerszten, Stephen B. Montgomery, Christopher B. Newgard, Eric A. Ortlund, Stuart C. Sealfon, Michael P. Snyder, Martin J. Walsh

**Clinical Sites:** Cheehoon Ahn, Alicia Belangee, Bryan C. Bergman, Daniel H. Bessesen, Gerard A. Boyd, Anna R. Brandt, Nicholas T. Broskey, Toby L. Chambers, Clarisa Chavez Martinez, Maria Chikina, Alex Claiborne, Zachary S. Clayton, Paul M. Coen, Katherine A. Collins-Bennett, Tiffany M. Cortes, Gary R. Cutter, Matthew Douglass, Daniel E. Forman, Will A. Fountain, Aaron H. Gouw, Kevin J. Gries, Fadia Haddad, Joseph A. Houmard, Kim M. Huffman, Ryan P. Hughes, John M. Jakicic, Catherine M. Jankowski, Neil M. Johannsen, Johanna L. Johnson, Erin E. Kershaw, Dillon J. Kuszmaul, Bridget Lester, Colleen E. Lynch, Edward L. Melanson, Cristhian Montenegro, Kerrie L. Moreau, Masatoshi Naruse, Jia Nie, Bradley C Nindl, Maria F. Pino, Tuomo Rankinen, Ulrika Raue, Ethan Robbins, Kaitlyn R. Rogers, Renee J. Rogers, Irene E. Schauer, Robert S. Schwartz, Chad M. Skiles, Lauren M. Sparks, Maja Stefanovic-Racic, Andrew M. Stroh, Kristen J. Sutton, Anna Thalacker-Mercer, Todd A. Trappe, Caroline S. Vincenty, Elena Volpi, Katie L. Whytock, Gilhyeon Yoon, Thomas W. Buford, Dan M. Cooper, Sara E. Espinoza, Bret H. Goodpaster, Wendy M. Kohrt, William E. Kraus, Nicolas Musi, Shlomit Radom-Aizik, Blake B. Rasmussen, Eric Ravussin, Scott Trappe

### MoTrPAC Study Group Acknowledgements

Nicole Adams, Abdalla Ahmed, Andrea Anderson, Carter Asef, Arianne Aslamy, Marcas M. Bamman, Jerry Barnes, Susan Barr, Kelsey Belski, Will Bennett, Amanda Boyce, Brandon Bukas, Emily Carifi, Chih-Yu Chen, Haiying Chen, Shyh-Huei Chen, Samuel Cohen, Audrey Collins, Gavin Connolly, Elaine Cornell, Julia Dauberger, Carola Ekelund, Shannon S. Emilson, Jerome Fleg, Nicole Gagne, Mary-Catherine George, Ellie Gibbons, Jillian Gillespie, Aditi Goyal, Bruce Graham, Xueyun Gulbin, Jere Hamilton, Leora Henkin, Andrew Hepler, Lidija Ivic, Ronald Jackson, Andrew Jones, Lyndon Joseph, Leslie Kelly, Gary Lee, Adrian Loubriel, Ching-ju Lu, Kristal M. Maner-Smith, Ryan Martin, Padma Maruvada, Alyssa Mathews, Curtis McGinity, Lucas Medsker, Kiril Minchev, Samuel G. Moore, Michael Muehlbauer, Anne Newman, John Nichols, Concepcion R. Nierras, George Papanicolaou, Lorrie Penry, June Pierce, Megan Reaves, Eric W. Reynolds, Teresa N. Richardson, Jeremy Rogers, Scott Rushing, Santiago Saldana, Rohan Shah, Samiya M. Shimly, Cris Slentz, Deanna Spaw, Debbie Steinberg, Suchitra Sudarshan, Alyssa Sudnick, Jennifer W. Talton, Christy Tebsherani, Nevyana Todorova, Jennifer Walker, Michael P. Walkup, Anthony Weakland, Gary Weaver, Christopher Webb, Sawyer Welden, John P. Williams, Marilyn Williams, Leslie Willis, Yi Zhang, Frank Booth, Andrea Hevener, Ian Lanza, Jun Li, K Sreekumaran Nair

## DECLARATION OF INTERESTS

The authors declare no competing interests.

## STAR Methods

### Experimental model and study participant details

#### IRB

The Molecular Transducers of Physical Activity Consortium (MoTrPAC) (NCT03960827) is a multicenter study designed to isolate the effects of structured exercise training on the molecular mechanisms underlying the health benefits of exercise and physical activity. Methods are described here in sufficient detail to interpret results, with references to supplemental material and prior publications for detailed descriptions. The present work includes data from prior to the suspension of the study in March 2020 due to the Covid-19 pandemic. The study was conducted in accordance with the Declaration of Helsinki and approved by the Institutional Review Board of Johns Hopkins University School of Medicine (IRB protocol # JHMIRB5; approval date: 05/06/2019). All MoTrPAC participants provided written informed consent for the MoTrPAC study indicating whether they agreed to open data sharing and at what level of sharing they wanted. Participants could choose to openly share all de-identified data, with the knowledge that they could be reidentified, and they could also choose to openly share limited de-identified individual level data, which is lower risk of re-identification. All analyses and resulting data and results are shared in compliance with the NIH Genomic Data Sharing (GDS) policy and DSMB requirements for the randomized study.

#### Participant Characteristics

Volunteers were screened to: (a) ensure they met all eligibility criteria; (b) acquire phenotypic assessments of the study population; and (c) verify they were healthy and medically cleared to be formally randomized into the study (cite Clinical Landscape).34 Participants were then randomized into intervention groups stratified by clinical site and completed a pre-intervention baseline acute test. A total of 176 participants completed the baseline acute test (EE=66, RE=73, and CON=37), and among those, 175 (99%) had at least one biospecimen sample collected. Participants then began 12 weeks of exercise training or control conditions. Upon completion of the respective interventions, participants repeated phenotypic testing and the acute test including biospecimen sampling. Due to the COVID-19 suspension, only 45 participants (26%) completed the post-intervention follow-up acute exercise test (some with less than 12 weeks of training), with 44 completing the post-intervention biospecimen collections. See Clinical Landscape for further detail.

#### Participant Assessments

As described in Clinical Landscape (REF, MoTrPAC Clinical landscape, *in review*) and protocol paper^73^, prior to randomization, participant screening was completed via questionnaires, and measurements of anthropometrics, resting heart rate and blood pressure, fasted blood panel, cardiorespiratory fitness, muscular strength, and free-living activity and sedentary behavior. Cardiorespiratory fitness was assessed using a cardiopulmonary exercise test (CPET) on a cycle ergometer (Lode Excalibur Sport Lode BV, Groningen, The Netherlands) with indirect calorimetry. Quadriceps strength was determined by isometric knee extension of the right leg at a 60° knee angle using a suitable strength device^73^. Grip strength of the dominant hand was obtained using the Jamar Handheld Hydraulic Dynamometer (JLW Instruments, Chicago, IL). See Clinical Landscape (REF, MoTrPAC Clinical Landscape, *in review*) for participant assessment results.

### Acute Exercise Intervention

The pre-intervention baseline EE acute bouts were composed of three parts: (1) a 5min warm up at 50% of the estimated power output to elicit 65% VO2peak, (2) 40min cycling at ∼65% VO2peak, and (3) a 1min recovery at ∼25 W. The pre-intervention baseline RE acute bouts were composed of a 5min warm-up at 50-60% heart rate reserve (HRR) followed by completion of three sets each of five upper (chest press, overhead press, seated row, triceps extension, biceps curl) and three lower body (leg press, leg curl, leg extension) exercises to volitional fatigue (∼10RM) with 90sec rest between each set. Participants randomized to CON did not complete exercise during their acute tests. Participants rested supine for 40min to mirror the EE and RE acute test schedule. See Clinical Landscape (REF, MoTrPAC Clinical Landscape, *in review*) for acute bout results.

### Adipose Tissue Collection and Omics Profiling

Abdominal subcutaneous adipose tissue (ASAT) biopsies were collected from all participants as part of the acute exercise test. Prior to testing, participants adhered to pre-sampling restrictions, including abstention from caffeine, alcohol, NSAIDs, exercise, and specific dietary components^73^. Upon arrival, participants rested supine for at least 30 minutes prior to the pre-exercise biopsy. Post-exercise adipose collections were conducted according to randomized temporal profiles designed to span the acute recovery period while minimizing participant burden. These profiles included early (Pre and 45 min Post), middle (Pre and 4 hr Post), late (Pre and 24 hr Post), and full (all four time points, although ASAT was only collected at Pre and 4 hr in this group). Final sample sizes were 43 for early, 80 for middle, and 49 for late profiles. The post-exercise biopsy was timed immediately following completion of the final exercise component—40 minutes of cycling for EE, the third set of leg extensions for RE, or the 40-minute rest period for CON. ASAT biopsies were successfully collected in 172 participants (EE = 62, RE = 73, CON = 37), and only these participants were included in this study.

Adipose tissue was subjected to multi-omics profiling. Transcriptomics and metabolomics analyses were performed at all four time points (Pre, 45 min, 4 hr, and 24 hr post-exercise). Proteomics and phosphoproteomics were conducted at Pre and 4 hr Post only. All biospecimen processing and quality control procedures followed standardized protocols as described in the Clinical Landscape (*REF: MoTrPAC Clinical Landscape, in review*) and protocol design manuscript^73^.

### Molecular Assays

#### Transcriptomics

RNA Sequencing (RNA-Seq) was performed at Stanford University and the Icahn School of Medicine at Mount Sinai.

##### Extraction of total RNA

Approximately 50 mg of frozen ASAT were disrupted in Agencourt RNAdvance tissue lysis buffer (Beckman Coulter, Brea, CA) using a tissue ruptor (Omni International, Kennesaw, GA, #19-040E). Total RNA was extracted in a BiomekFX automation workstation according to the manufacturer’s instructions for tissue-specific extraction. The RNA was quantified by NanoDrop (ThermoFisher Scientific, # ND-ONE-W) and Qubit assay (ThermoFisher Scientific), and the quality was determined by either Bioanalyzer or Fragment Analyzer analysis.

##### mRNA Sequencing Library Preparation

Universal Plus mRNA-Seq kit from NuGEN/Tecan (# 9133) were used for generation of RNA-Seq libraries derived from poly(A)-selected RNA according to the manufacturer’s instructions. Universal Plus mRNA-Seq libraries contain dual (i7 and i5) 8 bp barcodes and an 8 bp unique molecular identifier (UMI), which enable deep multiplexing of NGS sequencing samples and accurate quantification of PCR duplication levels. Approximately 300ng of ASAT was used. The Universal Plus mRNA-Seq workflow consists of poly(A) RNA selection, RNA fragmentation and double-stranded cDNA generation using a mixture of random and oligo(dT) priming, end repair to generate blunt ends, adaptor ligation, strand selection, AnyDeplete workflow to remove unwanted ribosomal and globin transcripts, and PCR amplification to enrich final library species. All library preparations were performed using a Biomek i7 laboratory automation system (Beckman Coulter). Tissue-specific reference standards provided by the consortium were included with all RNA isolations to QC the RNA.

##### RNA Sequencing, quantification, and normalization

RNA sequencing, quantification, and normalization Pooled libraries were sequenced on an Illumina NovaSeq 6000 platform (Illumina, San Diego, CA, USA) to a target depth of 40 million read pairs (80 million paired-end reads) per sample using a paired-end 100 base pair run configuration. In order to capture the 8-base UMIs, libraries were sequenced using 16 cycles for the i7 index read and 8 cycles for the i5 index read. Reads were demultiplexed with bcl2fastq2 (v2.20.0) using options --use-bases-mask Y*,I8Y*,I*,Y* --mask-short-adapter-reads 0 --minimum-trimmed-read-length 0 (Illumina, San Diego, CA, USA), and UMIs in the index FASTQ files were attached to the read FASTQ files. Adapters were trimmed with cutadapt (v1.18)^74^, and trimmed reads shorter than 20 base pairs were removed. FastQC (v0.11.8) was used to generate pre-alignment QC metrics. STAR (v2.7.0d)^75^ was used to index and align reads to release 38 of the Ensembl Homo sapiens (hg38) genome and Gencode (Version 29). Default parameters were used for STAR’s genomeGenerate run mode; in STAR’s alignReads run mode, SAM attributes were specified as NH HI AS NM MD nM, and reads were removed if they did not contain high-confidence collapsed splice junctions (--outFilterType BySJout). RSEM (v1.3.1)^76^ was used to quantify transcriptome-coordinate-sorted alignments using a forward probability of 0.5 to indicate a non-strand-specific protocol. Bowtie 2 (v2.3.4.3)^77^ was used to index and align reads to globin, rRNA, and phix sequences in order to quantify the percent of reads that mapped to these contaminants and spike-ins. UCSC’s gtfToGenePred was used to convert the hg38 gene annotation (GTF) to a refFlat file in order to run Picard CollectRnaSeqMetrics (v2.18.16) with options MINIMUM_LENGTH=50 and RRNA_FRAGMENT_PERCENTAGE=0.3. UMIs were used to accurately quantify PCR duplicates with NuGEN’s “nodup.py” script (https://github.com/tecangenomics/nudup). QC metrics from every stage of the quantification pipeline were compiled, in part with multiQC (v1.6). The openWDL-based implementation of the RNA-Seq pipeline on Google Cloud Platform is available on Github (https://github.com/MoTrPAC/motrpac-rna-seq-pipeline). Filtering of lowly expressed genes and normalization were performed separately in each tissue. RSEM gene counts were used to remove lowly expressed genes, defined as having 0.5 or fewer counts per million in at least 10% of samples. These filtered raw counts were used as input for differential analysis with the variancePartition::dream^78^, as described in the statistical analysis methods section. To generate normalized sample-level data for downstream visualization, filtered gene counts were TMM-normalized using edgeR::calcNormFactors, followed by conversion to log counts per million with edgeR::cpm.

Principal Component Analysis and calculation of the variance explained by variables of interest were used to identify and quantify potential batch effects. Based on this analysis, processing batch, clinical site, percentage of UMI duplication, and RNA Integrity Number (RIN) technical effects were regressed out of the TMM-normalized counts via linear regression using limma::RemoveBatchEffect^79^ function in R. A design matrix including age, sex, and a combination of group and timepoint was used during batch effect removal to avoid removing variance attributable to biological effects.

ASAT RNA extraction was performed in separate batches according to exercise modality, so the extraction batch and exercise modality were perfectly collinear. This collinearity was identified at the RNA extraction step and samples were randomized prior to construction of cDNA libraries. Ultimately, under this processing implementation, differences in gene expression attributable to exercise group can be impossible to disentangle from RNA extraction batch effects, so the batch variable was not regressed out in the technical effect stage or included as a covariate in the differential analysis, and left as an experimental limitation.

##### RNA Quality Inclusion Criteria

In the blood transcriptomic data, 79/1032 processed adult-sedentary samples had RIN values under 5. Based on established guidelines and internal quality control assessments, any samples with a RIN score below 5 were excluded from further analysis due to concerns about potential degradation artifacts. Additional visualizations and summary figures supporting this decision are available in the quality control report at: https://github.com/MoTrPAC/precovid-analyses/tree/main/QC/

### Proteomics and Phosphoproteomics

#### Study design

LC-MS/MS analysis was performed at the Pacific Northwest National Laboratories (PNNL). 46 ASAT samples representing baseline and 4-hours post-intervention from all three groups were analyzed.

#### Generation of common reference

Samples were digested at each site following sample processing protocol described below, and centrally aliquoted. Common reference for adipose tissue was generated using bulk tissue from 6 individuals representing a 4:2 ratio of female:male. 250 μg aliquots of common reference samples were made to be included in each multiplex (described below) and additional aliquots are stored for inclusion in future MoTrPAC phases to facilitate data integration.

#### Sample processing

Proteomics analyses were performed using clinical proteomics protocols described previously^80,81^. Samples were lysed in ice-cold, freshly-prepared lysis buffer (8 M urea (Sigma-Aldrich, St. Louis, Missouri), 50 mM Tris pH 8.0, 75 mM sodium chloride, 1 mM EDTA, 2 μg/ml Aprotinin (Sigma-Aldrich, St. Louis, Missouri), 10 μg/ml Leupeptin (Roche CustomBiotech, Indianapolis, Indiana), 1 mM PMSF in EtOH, 10 mM sodium fluoride, 1% phosphatase inhibitor cocktail 2 and 3 (Sigma-Aldrich, St. Louis, Missouri), 10 mM Sodium Butyrate, 2 μM SAHA, and 10 mM nicotinamide and protein concentration was determined by BCA assay. Protein lysate concentrations were normalized within samples of the same tissue type, and protein was reduced with 5 mM dithiothreitol (DTT, Sigma-Aldrich) for 1 hour at 37°C with shaking at 1000 rpm on a thermomixer, alkylated with iodoacetamide (IAA, Sigma-Aldrich) in the dark for 45 minutes at 25°C with shaking at 1000 rpm, followed by dilution of 1:4 with Tris-HCl, pH 8.0 prior to adding digestion enzymes. Proteins were first digested with LysC endopeptidase (Wako Chemicals) at a 1:50 enzyme:substrate ratio (2 hours, 25 °C, 850 rpm), followed by digestion with trypsin (Promega) at a 1:50 enzyme:substrate ratio (or 1:10 ratio for adipose tissue; 14 hours, 25 °C, 850 rpm). The next day formic acid was added to a final concentration of 1% to quench the reaction. Digested peptides were desalted using Sep-Pac C18 columns (Waters), concentrated in a vacuum centrifuge, and a BCA assay was used to determine final peptide concentrations. 250μg aliquots of each sample were prepared, dried down by vacuum centrifugation and stored at -80°C.

Tandem mass tag (TMT) 16-plex isobaric labeling reagent (ThermoFisher Scientific) was used for this study. Samples were randomized across the first 15 channels of TMT 16-plexes, and the last channel (134N) of each multiplex was used for a common reference that was prepared prior to starting the study (see above). Randomization of samples across the plexes within each site was done using https://github.com/MoTrPAC/clinical-sample-batching, with the goal to have all timepoints per participant in the same plex, and uniform distribution of groups (endurance, resistance, control), sex and sample collection clinical site across the plexes. Peptide aliquots (250 μg per sample) were resuspended to a final concentration of 5 μg/μL in 200 mM HEPES, pH 8.5 for isobaric labeling. TMT reagent was added to each sample at a 1:2 peptide: TMT ratio, and labeling proceeded for 1 hour at 25°C with shaking at 400 rpm. The labeling reaction was diluted to a peptide concentration of 2 µg/µL using 62.5 μL of 200 mM HEPES and 20% ACN. 3 μL was removed from each sample to quantify labeling efficiency and mixing ratio. After labeling QC analysis, reactions were quenched with 5% hydroxylamine and samples within each multiplex were combined and desalted with Sep-Pac C18 columns (Waters).

Combined TMT multiplexed samples were then fractionated using high pH reversed phase chromatography on a 4.6mm ID x 250mm length Zorbax 300 Extend-C18 column (Agilent) with 5% ammonium formate/2% Acetonitrile as solvent A and 5% ammonium formate/90% acetonitrile as solvent B. Samples were fractionated with 96min separation gradient at flow rate of 1mL/min and fractions were collected at each minute onto a 96-well plate. Fractions are then concatenated into 24 fractions with the following scheme: fraction 1 = A1+C1+E1+G1, fraction 2 = A2+C2+E2+G2, fraction 3 = A3+C3+E3+G3, all the way to fraction 24 = B12+D12+F12+H12 following the same scheme. 5% of each fraction was removed for global proteome analysis, and the remaining 95% was further concatenated to 12 fractions for phosphopeptide enrichment using immobilized metal affinity chromatography (IMAC).

Phosphopeptide enrichment was performed through immobilized metal affinity chromatography (IMAC) using Fe 3+ -NTA-agarose beads, freshly prepared from Ni-NTA-agarose beads (Qiagen, Hilden, Germany) by sequential incubation in 100 mM EDTA to strip nickel, washing with HPLC water, and incubation in 10 mM iron (III) chloride). Peptide fractions were resuspended to 0.5 μg/uL in 80% ACN + 0.1% TFA and incubated with beads for 30 minutes in a thermomixer set to 1000 rpm at room temperature. After 30 minutes, beads were spun down (1 minute, 1000 rcf) and supernatant was removed and saved as flow-through for subsequent enrichments. Phosphopeptides were eluted off IMAC beads in 3x 75 μL of agarose bead elution buffer (500 mM K2HPO4, pH 7.0), desalted using C18 stage tips, eluted with 50% ACN, and lyophilized. Samples were then reconstituted in 3% ACN / 0.1% FA for LC-MS/MS analysis (9 μL reconstitution / 4 μL injection at the BI; 12 μL reconstitution / 5 μL injection at PNNL).

#### Data acquisition

For mass spectrometry analysis of the global proteome, online separation was performed using a Dionex Ultimate 3000 UHPLC system (ThermoFisher) and a 25 cm x 75 μm i.d. picofrit column packed in-house with C18 silica (1.7 μm UPLC BEH particles, Waters Acquity) with solvent A of 0.1% formic acid/3% acetonitrile and solvent B of 0.1% formic acid/90% acetonitrile, flow rate of 200nL/min and the following gradient: 1-8% B in 10min, 8-25% B in 90min, 25-35% B in 10min, 35-75% B in 5min, and 75-5% in 3min. Proteome fractions were analyzed on a Q Exactive HF-X Plus mass spectrometer (ThermoFisher Scientific) with MS1 scan across the 300-1800 m/z range at 60,000 resolution, AGC target of 3×106 and maximum injection time of 20ms. MS2 scans of most abundant 12 ions were performed at 30,000 resolution with AGC target of 1×105 and maximum injection time of 100ms, isolation window of 0.7m/z and normalized collision energy of 30.

For mass spectrometry analysis of the phosphoproteome, online separation was performed using a Dionex Ultimate 3000 UHPLC system (ThermoFisher) and a 25 cm x 75 μm i.d. picofrit column packed in-house with C18 silica (1.7 μm UPLC BEH particles, Waters Acquity) with solvent A of 0.1% formic acid/3% acetonitrile and solvent B of 0.1% formic acid/90% acetonitrile, flow rate of 200nL/min and the following gradient: 1-8% B in 10min, 8-25% B in 90min, 25-35% B in 10min, 35-75% B in 5min, and 75-5% in 3min. Phosphoproteome fractions were analyzed on a Q Exactive HF-X Plus mass spectrometer (ThermoFisher Scientific) with MS1 scan across the 300-1800 m/z range at 60,000 resolution, AGC target of 3×106 and maximum injection time of 20ms. MS2 scans of most abundant 12 ions were performed at 45,000 resolution with AGC target of 1×105 and maximum injection time of 100ms, isolation window of 0.7m/z and normalized collision energy of 30.

#### Data searching

Proteome and phosphoproteome data were searched against a composite protein database at the Bioinformatics Center (BIC) using the MSGF+ cloud-based pipeline previously described^82^. This database comprised UniProt canonical sequences (downloaded 2022-09-13; 20383 sequences), UniProt human protein isoforms (downloaded 2022-09-13; 21982 sequences), and common contaminants (261 sequences), resulting in 42,626 sequences.

#### QC/Filtering/Normalization

The log2 Reporter ion intensity (RII) ratios to the common reference were used as quantitative values for all proteomics features (proteins and phosphosites). Datasets were filtered to remove features identified from contaminant proteins and decoy sequences. Datasets were visually evaluated for sample outliers by looking at top principal components, examining median feature abundance and distributions of RII ratio values across samples, and by quantifying the number of feature identifications within each sample. No outliers were detected in the muscle tissue dataset. In the adipose datasets (proteome and phosphoproteome), four samples were flagged as outliers based on inspection of the top principal components and evaluation of the raw quantitative results (median feature abundance <-1.0). The Log2 RII ratio values were normalized within each sample by median centering to zero. Principal Component Analysis and calculation of the variance explained by variables of interest were used to identify and quantify potential batch effects. Based on this analysis, TMT plex, and Clinical Site batch effects were removed using Linear Models Implemented in the limma::RemoveBatchEffect() function in R^79^. A design matrix including age, sex, and group_timepoint was used during batch effect removal in order to preserve the effect of all variables included in later statistical analysis. Correlations between technical replicates analyzed within and across CAS (where applicable) were calculated to evaluate intra- and inter-site reproducibility; the data from technical replicates were then averaged for downstream analysis. Finally, features with quantification in less of 30% of all samples were removed. For specific details of the process, see available code.

### Metabolomics and Lipidomics

The metabolomic analysis was performed by investigators across multiple Chemical Analysis Sites that employed different technical platforms for data acquisition. At the highest level, these platforms were divided into two classes: Targeted and Untargeted. The data generated by the untargeted platforms were further divided into Named (confidently identified chemical entities) and Unnamed (confidently detected, but no chemical name annotation) compounds.

Targeted metabolomics data were generated at 3 analysis Sites: Duke University, the Mayo Clinic, and Emory University. Duke quantified metabolites belonging to the metabolite classes Acetyl-CoA, Keto Acids, and Nucleic Acids (“acoa”, “ka”, “nuc”), Mayo quantified amines and TCA intermediates (“amines” and “tca”), while Emory quantified Oxylipins (“oxylipneg”).

The untargeted metabolomics data were generated at 3 analysis Sites: the Broad Institute, the University of Michigan, and the Georgia Institute of Technology (Georgia Tech). The Broad Institute applied Hydrophilic interaction liquid chromatography (HILIC) in the positive ion mode (hilicpos), Michigan applied reverse phase liquid chromatography in both positive and negative ion modes (“rppos” and “rpneg”) and ion-pairing chromatography in the negative mode (“ionpneg”), and Georgia Tech performed lipidomics assays using reverse phase chromatography in both positive and negative ion modes (“lrppos” and “lrpneg”).

#### LC-MS/MS analysis of branched-chain keto acids

Duke University conducted targeted profiling of branched-chain keto acid metabolites. Plasma samples containing isotopically labeled ketoleucine (KIC)-d3, ketoisovalerate (KIV)-13C5 (from Cambridge Isotope Laboratories), and 3-methyl-2-oxovalerate (KMV)-d8 (from Toronto Research Chemicals, Canada) internal standards, were subjected to deproteinization with 3M perchloric acid. Tissue homogenates were prepared at 100 mg/ml in 3M perchloric acid and 200 μL was centrifuged.

Next, 200 μL of 25 M o-phenylenediamine (OPD) in 3M HCl were added to both the plasma and tissue supernatants. The samples were incubated at 80°C for 20 minutes. Keto acids were then extracted using ethyl acetate, following a previously described protocol^83,84^. The extracts were dried under nitrogen, reconstituted in 200 mM ammonium acetate, and subjected to analysis on a Xevo TQ-S triple quadrupole mass spectrometer coupled to an Acquity UPLC (Waters), controlled by the MassLynx 4.1 operating system.

The analytical column used was a Waters Acquity UPLC BEH C18 Column (1.7 μm, 2.1 × 50 mm), maintained at 30°C. 10 μL of the sample were injected onto the column and eluted at a flow rate of 0.4 ml/min. The gradient consisted of 45% mobile phase A (5 mM ammonium acetate in water) and 55% mobile phase B (methanol) for 2 minutes. This was succeeded by a linear gradient to 95% B from 2 to 2.5 minutes, holding at 95% B for 0.7 minutes, returning to 45% A, and finally re-equilibrating the column at initial conditions for 1 minute. The total run time was 4.7 minutes.

In positive ion mode, mass transitions of m/z 203 → 161 (KIC), 206 → 161 (KIC-d3), 189 → 174 (KIV), 194 → 178 (KIV-5C13), 203 → 174 (KMV), and 211 → 177 (KMV-d8) were monitored. Endogenous keto acids were quantified using calibrators prepared by spiking dialyzed fetal bovine serum with authentic keto acids (Sigma-Aldrich).

#### Flow injection MS/MS analysis of acyl CoAs

Duke University conducted targeted profiling of acyl CoAs. Here, 500 μL of tissue homogenate, prepared at a concentration of 50 mg/ml in isopropanol/0.1 M KH2PO4 (1:1), underwent extraction with an equal volume of acetonitrile. The resulting mixture was then centrifuged at 14,000 x g for 10 minutes, following a previously established procedure^85,86^.

The supernatants were acidified with 0.25 ml of glacial acetic acid, and the acyl CoAs were subjected to additional purification through solid-phase extraction (SPE) using 2-(2-pyridyl) ethyl functionalized silica gel (Sigma-Aldrich), as outlined in Minkler et al. (2008)^87^. Prior to use, the SPE columns were conditioned with 1 ml of acetonitrile/isopropanol/water/glacial acetic acid (9/3/4/4 : v/v/v/v). After application and flow-through of the supernatant, the SPE columns underwent a washing step with 2 ml of acetonitrile/isopropanol/water/glacial acetic acid (9/3/4/4 : v/v/v/v). Acyl CoAs elution was achieved with 2 ml of methanol/250 mM ammonium formate (4/1 : v/v), followed by analysis using flow injection MS/MS in positive ion mode on a Xevo TQ-S triple quadrupole mass spectrometer (Waters). The mobile phase used was methanol/water (80/20, v/v) with 30 mM ammonium hydroxide. Spectra were acquired in the multichannel acquisition mode, monitoring the neutral loss of 507 amu (phosphoadenosine diphosphate) and scanning from m/z 750 to 1100. As an internal standard, heptadecanoyl CoA was employed. Quantification of endogenous acyl CoAs was carried out using calibrators created by spiking tissue homogenates with authentic Acyl CoAs (Sigma-Aldrich) that covered saturated acyl chain lengths from C0 to C18. Empirical corrections for heavy isotope effects, particularly 13C, to the adjacent m+2 spectral peaks in a specific chain length cluster were made by referring to the observed spectra for the analytical standards.

#### LC-MS/MS analysis of nucleotides

Duke University conducted targeted profiling of nucleotide metabolites. 300 μL of tissue homogenates, prepared at a concentration of 50 mg/ml in 70% methanol, underwent spiking with nine internal standards: 13C10,15N5-adenosine monophosphate, 13C10,15N5-guanosine monophosphate, 13C10,15N2-uridine monophosphate, 13C9,15N3-cytidine monophosphate, 13C10-guanosine triphosphate, 13C10-uridine triphosphate, 13C9-cytidine triphosphate, 13C10-adenosine triphosphate, and nicotinamide-1, N6-ethenoadenine dinucleotide (eNAD) (all from Sigma-Aldrich). Nucleotides were extracted using an equal volume of hexane, following the procedures previously outlined^88,89^. After vortexing and centrifugation at 14,000 x g for 5 minutes, the bottom layer was subjected to another centrifugation. Chromatographic separation and MS analysis of the supernatants were performed using an Acquity UPLC system (Waters) coupled to a Xevo TQ-XS quadrupole mass spectrometer (Waters). The analytical column employed was a Chromolith FastGradient RP-18e 50-2mm column (EMD Millipore, Billerica, MA, USA), that was maintained at 40°C. The injection volume was 2 μL. Nucleotides were separated using a mobile phase A consisting of 95% water, 5% methanol, and 5 mM dimethylhexylamine adjusted to pH 7.5 with acetic acid. Mobile phase B comprised 20% water, 80% methanol, and 10 mM dimethylhexylamine. The flow rate was set to 0.3 ml/min. The 22-minute gradient (t=0, %B=0; t=1.2, %B=0; t=22, %B=40) was followed by a 3-minute wash and 7-minute equilibration. Nucleotides were detected in the negative ion multiple reaction monitoring (MRM) mode based on characteristic fragmentation reactions. Endogenous nucleotides were quantified using calibrators prepared by spiked-in authentic nucleotides obtained from Sigma-Aldrich.

#### LC-MS/MS analysis of amino metabolites

The Mayo Clinic conducted LC-MS-based targeted profiling of amino acids and amino metabolites, as previously outlined^90,91^. In summary, either 20 ml of plasma samples or 5 mg of tissue homogenates were supplemented with an internal standard solution comprising isotopically labeled amino acids (U-13C4 L-aspartic acid, U-13C3 alanine, U-13C4 L-threonine, U-13C L-proline, U-13C6 tyrosine, U-13C5 valine, U-13C6 leucine, U-13C6 phenylalanine, U-13C3 serine, U-13C5 glutamine, U-13C2 glycine, U-13C5 glutamate, U-13C6,15N2 lysine, U-13C5,15N methionine, 1,1U-13C2 homocysteine, U-13C6 arginine, U-13C5 ornithine, 13C4 asparagine, 13C2 ethanolamine, D3 sarcosine, D6 4-aminobutyric acid). The supernatant was promptly derivatized using 6-aminoquinolyl-N-hydroxysuccinimidyl carbamate with a MassTrak kit (Waters). A 10-point calibration standard curve underwent a similar derivatization procedure after the addition of internal standards. Both the derivatized standards and samples were subjected to analysis using a Quantum Ultra triple quadrupole mass spectrometer (ThermoFischer) coupled with an Acquity liquid chromatography system (Waters). Data acquisition utilized selected ion monitoring (SRM) in positive ion mode. The concentrations of 42 analytes in each unknown sample were calculated against their respective calibration curves.

#### GC-MS analysis of TCA metabolites

The Mayo Clinic conducted GC-MS-targeted profiling of tricarboxylic acid (TCA) metabolites, as previously detailed^92,93^, with some modifications. In summary, 5 mg of tissue were homogenized in 1X PBS using an Omni bead ruptor (Omni International, Kennesaw, GA), followed by the addition of 20 μL of an internal solution containing U-13C labeled analytes (13C3 sodium lactate, 13C4 succinic acid, 13C4 fumaric acid, 13C4 alpha-ketoglutaric acid, 13C4 malic acid, 13C4 aspartic acid, 13C5 2-hydroxyglutaric acid, 13C5 glutamic acid, 13C6 citric acid, 13C2,15N glycine, 13C2 sodium pyruvate). For plasma, 50 μL were used. The proteins were precipitated out using 300 μL of a mixture of chilled methanol and acetonitrile solution. After drying the supernatant in the speedvac, the sample was derivatized first with ethoxime and then with MtBSTFA + 1% tBDMCS (N-Methyl-N-(t-Butyldimethylsilyl)-Trifluoroacetamide + 1% t-Butyldimethylchlorosilane) and then analyzed on an Agilent 5977B GC/MS (Santa Clara, California) under single ion monitoring conditions using electron ionization. Concentrations of lactic acid (m/z 261.2), fumaric acid (m/z 287.1), succinic acid (m/z 289.1), ketoglutaric acid (m/z 360.2), malic acid (m/z 419.3), aspartic acid (m/z 418.2), 2-hydroxyglutaratic acid (m/z 433.2), cis-aconitic acid (m/z 459.3), citric acid (m/z 591.4), isocitric acid (m/z 591.4), and glutamic acid (m/z 432.4) were measured against 7-point calibration curves that underwent the same derivatization procedure.

#### Targeted lipidomics of low-level lipids

Lipid targeted profiling was conducted at Emory University following established methodologies as previously described^94,95^. In summary, 20 mg of powdered ASAT samples were homogenized in 100 μL PBS using Bead Ruptor (Omni International, Kennesaw, GA). Homogenized tissue samples (or plasma samples) were diluted with 300 μL 20% methanol and spiked with a 1% BHT solution to a final BHT concentration of 0.1% and pH of 3.0 by acetic acid addition. After centrifugation (10 minutes, 14000 rpm), the supernatants were transferred to 96-well plates for further extraction. The supernatants were loaded onto Isolute C18 solid-phase extraction (SPE) columns that had been conditioned with 1000 μL ethyl acetate and 1000 μL 5% methanol. The SPE columns were washed with 800 μL water and 800 μL hexane, and the oxylipins were then eluted with 400 μL methyl formate. The SPE process was automated using a Biotage Extrahera (Uppsala, Sweden). The eluate was dried with nitrogen and reconstituted with 200 μL acetonitrile before LC-MS analysis. Sample blanks, pooled extract samples used as quality controls (QC), and consortium reference samples were prepared for analysis using the same methods. All external standards were purchased from Cayman Chemical (Ann Arbor, Michigan) at a final concentration in the range 0.01-20 μg/ml and consisted of: prostaglandin E2 ethanolamide (Catalog No. 100007212), oleoyl ethanolamide (Catalog No. 90265), palmitoyl ethanolamide (Catalog No. 10965), arachidonoyl ethanolamide (Catalog No. 1007270), docosahexaenoyl ethanolamide (Catalog No. 10007534), linoleoyl ethanolamide (Catalog No. 90155), stearoyl ethanolamide (Catalog No. 90245), oxy-arachidonoyl ethanolamide (Catalog No. 10008642), 2-arachidonyl glycerol (Catalog No. 62160), docosatetraenoyl ethanolamide (Catalog No. 90215), α-linolenoyl ethanolamide (Catalog No. 902150), oleamide (Catalog No. 90375), dihomo-γ-linolenoyl ethanolamide (Catalog No. 09235), decosanoyl ethanolamide (Catalog No. 10005823), 9,10 DiHOME (Catalog No. 53400), prostaglandin E2-1-glyceryl ester (Catalog No. 14010), 20-HETE (Catalog No. 10007269), 9-HETE (Catalog No. 34400), 14,15 DiHET (Catalog No. 10007267), 5(S)-HETE (Catalog No. 34210), 12(R)-HETE (Catalog No. 10007247), 11(12)-DiHET (Catalog No. 10007266), 5,6-DiHET (Catalog No. 10007264), thromboxane B2 (Catalog No. 10007237), 12(13)-EpOME (Catalog No. 52450), 13 HODE (Catalog No. 38600), prostaglandin F2α (Catalog No. 10007221), 14(15)-EET (Catalog No. 10007263), 8(9)-EET (Catalog No. 10007261), 11(12)-EET (Catalog No. 10007262), leukotriene B4 (Catalog No. 20110), 8(9)-DiHET (Catalog No. 10007265), 13-OxoODE (Catalog No. 38620), 13(S)-HpODE (Catalog No. 48610), 9(S)-HpODE (Catalog No. 48410), 9(S)-HODE (Catalog No. 38410), resolvin D3 (Catalog No. 13834), resolvin E1 (Catalog No. 10007848), resolvin D1 (Catalog No. 10012554), resolvin D2 (Catalog No. 10007279), 9(S)HOTrE (Catalog No. 39420), 13(S)HOTrE (Catalog No. 39620), 8-iso Progstaglandin F2α (Catalog No. 25903), maresin 1 (Catalog No. 10878), maresin 2 (Catalog No. 16369).

LC-MS data were acquired using an Agilent 1290 Infinity II chromatograph from Agilent (Santa Clara, CA), equipped with a ThermoFisher Scientific AccucoreTM C18 column (100 mm × 4.6, 2.6 µm particle size), and coupled to Agilent 6495 mass spectrometer for polarity switch multiple reaction monitoring (MRM) scan. The mobile phases consisted of water with 10 mM ammonium acetate (mobile phase A) and acetonitrile with 10 mM ammonium acetate (mobile phase B). The chromatographic gradient program was: 0.5 minutes with 95% A; 1 minute to 2 minutes with 65% A; 2.1 minutes to 5.0 minutes with 45% A; 7 minutes to 20 minutes with 25% A; and 21.1 minutes until 25 minutes with 95% A. The flow rate was set at 0.40 ml/min. The column temperature was maintained at 35°C, and the injection volume was 6 μL. For MS analysis, the MRM analysis is operated at gas temperature of 290 °C, gas flow of 14 L/min, Nebulizer of 20 psi, sheath gas temperature of 300 °C, sheath gas flow of 11 L /min, capillary of 3000 V for both positive and negative ion mode, nozzle voltage of 1500 V for both positive and negative ion mode, high pressure RF of iFunnel parameters of 150V for both positive and negative ion mode, and low pressure of RF of iFunnel parameters of 60 V for both positive and negative ion mode. Skyline (version 25.1.0.142)154 was utilized to process raw LC-MS data. Standard curves were constructed for each oxylipin/ethanolamide and scrutinized to ensure that all concentration points fell within the linear portion of the curve with an R-squared value not less than 0.9. Additionally, features exhibiting a high coefficient of variation (CV) among the quality control (QC) samples were eliminated from the dataset. Pearson correlation among the QCs for each tissue type was computed using the Hmisc R library, and the figures documented in the QC report were generated and visualized with the corrplot R library^96,97^.

#### Hydrophilic interaction LC-MS metabolomics

The untargeted analysis of polar metabolites in the positive ionization mode was conducted at the Broad Institute of MIT and Harvard. The LC-MS system consisted of a Shimadzu Nexera X2 UHPLC (Shimadzu Corp., Kyoto, Japan) coupled to a Q-Exactive hybrid quadrupole Orbitrap mass spectrometer (Thermo Fisher Scientific). Plasma samples (10 μL) were extracted using 90 μL of 74.9/24.9/0.2 v/v/v acetonitrile/methanol/formic acid containing valine-d8 and phenylalanine-d8 internal standards. Following centrifugation, the supernatants were directly injected onto a 150 x 2 mm, 3 µm Atlantis HILIC column (Waters). Tissue (10 mg) homogenization was performed at 4°C using a TissueLyser II (QAIGEN) bead mill set to two 2 min intervals at 20 Hz in 300 μL of 10/67.4/22.4/0.018 v/v/v/v water/acetonitrile/methanol/formic acid valine-d8 and phenylalanine-d8 internal standards. The column was eluted isocratically at a flow rate of 250 μL/min with 5% mobile phase A (10 mM ammonium formate and 0.1% formic acid in water) for 0.5 minute, followed by a linear gradient to 40% mobile phase B (acetonitrile with 0.1% formic acid) over 10 minutes, then held at 40% B for 4.5 minutes. MS analyses utilized electrospray ionization in the positive ion mode, employing full scan analysis over 70-800 m/z at 70,000 resolution and 3 Hz data acquisition rate. Various MS settings, including sheath gas, auxiliary gas, spray voltage, and others, were specified for optimal performance. Data quality assurance was performed by confirming LC-MS system performance with a mixture of >140 well-characterized synthetic reference compounds, daily evaluation of internal standard signals, and the analysis of four pairs of pooled extract samples per sample type inserted in the analysis queue at regular intervals. One sample from each pair was used to correct for instrument drift using “nearest neighbor” scaling while the second reference sample served as a passive QC for determination of the analytical coefficient of variation of every identified metabolite and unknown. Raw data processing involved the use of TraceFinder software (v3.3, Thermo Fisher Scientific) for targeted peak integration and manual review, as well as Progenesis QI (v3.0, Nonlinear Dynamics, Waters) for peak detection and integration of both identified and unknown metabolites. Metabolite identities were confirmed using authentic reference standards.

#### Reversed Phase-High Performance & Ion-pairing LC metabolomics

Reversed-phase and ion pairing LC-MS profiling of polar metabolites was conducted at the University of Michigan. LC-MS grade solvents and mobile phase additives were procured from Sigma-Aldrich, while chemical standards were obtained from either Sigma-Aldrich or Cambridge Isotope Labs. Plasma (50 μL aliquot) was extracted by addition of 200 μL of 1:1:1 v:v methanol:acetonitrile:acetone containing the following internal standards diluted from stock solutions to yield the specified concentrations: D4-succinic acid, 12.5 µM; :D3-malic acid,12.5 µM; D5 -glutamic acid,12.5 µM; D10 -leucine,12.5 µM; D5-tryptophan,12.5 µM;D5-phenylalanine,12.5 µM; D3-caffeine,12.5 µM; D8-lysine,12.5 µM; D4-chenodeoxycholic acid,12.5 µM; phosphatidylcholine(17:0/17:0),2.5 µM; phosphatidylcholine(19:0/19:0),2.5 µM; D31-palmitic acid,12.5 µM; D35-stearic acid,12.5 µM; gibberellic acid, 2.5 µM; epibrassinolide, 2.5 µM. The extraction solvent also contained a 1:400 dilution of Cambridge Isotope carnitine/acylcarnitine mix NSK-B, resulting in the following concentrations of carnitine/acylcarnitine internal standards: D9-L-carnitine, 380 nM; D3-L-acetylcarnitine, 95 nM; D3-L-propionylcarnitine, 19 nM; D3-L-butyrylcarnitine, 19 nM; D9-L-isovalerylcarnitine, 19 nM; D3-L-octanoylcarnitine, 19 nM; D9-L-myristoylcarnitine, 19 nM; D3-L-palmitoylcarnitine, 38 nM. After addition of solvent, samples were vortexed, incubated on ice for 10 minutes, and then centrifuged at 15,000 x g for 5 minutes. 150 μL supernatant were retrieved from the pellet, transferred to a glass autosampler vial with a 250 uL footed insert, and dried under a constant stream of nitrogen gas at ambient temperature. A QC sample was created by pooling residual supernatant from multiple samples. This QC sample underwent the same drying and reconstitution process as described for individual samples. Dried supernatants were stored at - 80 C until ready for instrumental analysis. On the day of analysis, dried samples were reconstituted in 37.5 μL of 8:2 v:v water:methanol, vortexed thoroughly, and submitted for LC-MS.

Frozen ASAT was rapidly weighed into pre-tared, pre-chilled Eppendorf tubes and extracted in 1:1:1:1 v:v methanol:acetonitrile:acetone:water containing the following internal standards diluted from stock solutions to yield the specified concentrations: 13C3-lactic acid, 12.5 µM; 13C5-oxoglutaric acid, 125 nM;13C5-citric acid,1.25 µM; 13C4-succinic acid,125 nM; 13C4-malic acid, 125 nM; U-13C amino acid mix (Cambridge Isotope CLM-1548-1), 2.5 µg/mL; 13C5-glutamine, 6.25 µM;15N2-asparagine, 1.25 µM;15N2-tryptophan, 1.25 µM; 13C6-glucose, 62.5 µM; D4-thymine, 1 µM; 15N-anthranillic acid, 1 µM; gibberellic acid,1 µM; epibrassinolide, 1 µM. The extraction solvent also contained a 1:400 dilution of Cambridge Isotope carnitine/acylcarnitine mix NSK-B, resulting in the following concentrations of carnitine/acylcarnitine internal standards: D9-L-carnitine, 380 nM; D3-L-acetylcarnitine, 95 nM; D3-L-propionylcarnitine, 19 nM; D3-L-butyrylcarnitine, 19 nM; D9-L-isovalerylcarnitine, 19 nM; D3-L-octanoylcarnitine, 19 nM; D9-L-myristoylcarnitine, 19 nM; D3-L-palmitoylcarnitine, 38 nM. Sample extraction was performed by adding chilled extraction solvent to tissue sample at a ratio of 1 ml solvent to 50 mg wet tissue mass. Immediately after solvent addition, the sample was homogenized using a Branson 450 probe sonicator. Subsequently, the tubes were mixed several times by inversion and then incubated on ice for 10 minutes. Following incubation, the samples were centrifuged and 300 μL of the supernatant was carefully transferred to two autosampler vials with flat-bottom inserts and dried under a constant stream of nitrogen gas. A QC sample was created by pooling residual supernatants from multiple samples. This QC sample underwent the same drying and reconstitution process as described for individual samples. Dried supernatants were stored at -80 C until ready for instrumental analysis. On the day of analysis, samples were reconstituted in 60 μL of 8:2 v:v water:methanol and then submitted to LC-MS.

Reversed phase LC-MS samples were analyzed on an Agilent 1290 Infinity II / 6545 qTOF MS system with a JetStream electrospray ionization (ESI) source (Agilent Technologies, Santa Clara, California) using a Waters Acquity HSS T3 column, 1.8 µm 2.1 x 100 mm equipped with a matched Vanguard precolumn (Waters Corporation). Mobile phase A was 100% water with 0.1% formic acid and mobile phase B was 100% methanol with 0.025% formic acid. The gradient was as follows: Linear ramp from 0% to 100% B from 0-10 minutes, hold 100% B until 17 minutes, linear return to 0% B from 17 to 17.1 minutes, hold 0% B until 20 minutes. The flow rate was 0.45 ml/min, the column temperature was 55°C, and the injection volume was 5 μL. Each sample was analyzed twice, once in positive and once in negative ion mode MS, scan rate 2 spectra/sec, mass range 50-1200 m/z. Source parameters were: drying gas temperature 350°C, drying gas flow rate 10 L/min, nebulizer pressure 30 psig, sheath gas temperature 350°C and flow 11 L/minute, capillary voltage 3500 V, internal reference mass correction enabled. A QC sample run was performed at minimum every tenth injection.

Ion-pair LC-MS samples were analyzed on an identically-configured LC-MS system using an Agilent Zorbax Extend C18 1.8 µm RRHD column, 2.1 x 150 mm ID, equipped with a matched guard column. Mobile phase A was 97% water, 3% methanol. Mobile phase B was 100% methanol. Both mobile phases contained 15 mM tributylamine and 10 mM acetic acid. Mobile phase C was 100% acetonitrile. Elution was carried out using a linear gradient followed by a multi-step column wash including automated (valve-controlled) backflushing, detailed as follows: 0-2min, 0%B; 2-11 min, linear ramp from 0-99%B; 12-16 min, 99%B, 16-17.5min, 99-0%B. At

17.55 minutes, the 10-port valve was switched to reverse flow (back-flush) through the column. From 17.55-20.45 min the solvent was ramped from 99%B to 99%C. From 20.45-20.95 min the flow rate was ramped up to 0.8 mL/min, which was held until 22.45 min, then ramped down to 0.6mL/min by 22.65 min. From 22.65-23.45 min the solvent was ramped from 99% to 0% C while flow was simultaneously ramped down from 0.6-0.4mL/min. From 23.45 to 29.35 min the flow was ramped from 0.4 to 0.25mL/min; the 10-port valve was returned to restore forward flow through the column at 25 min. . Column temperature was 35°C and the injection volume was 5 μL. MS acquisition was performed in negative ion mode, scan rate 2 spectra/sec, mass range 50-1200 m/z. Source parameters were: drying gas temperature 250°C, drying gas flow rate 13 L/min, nebulizer pressure 35 psig, sheath gas temp 325°C and flow 12 L/min, capillary voltage 3500V, internal reference mass correction enabled. A QC sample run was performed at minimum every tenth injection.

Iterative MS/MS data was acquired for both reverse phase and ion pairing methods using the pooled sample material to enable compound identification. Eight repeated LC-MS/MS runs of the QC sample were performed at three different collision energies (10, 20, and 40) with iterative MS/MS acquisition enabled. The software excluded precursor ions from MS/MS acquisition within 0.5 minute of their MS/MS acquisition time in prior runs, resulting in deeper MS/MS coverage of lower-abundance precursor ions.

Feature detection and alignment was performed utilizing a hybrid targeted/untargeted approach. Targeted compound detection and relative quantitation was performed by automatic integration followed by manual inspection and correction using Profinder v8.0 (Agilent Technologies, Santa Clara, CA.) Non-targeted feature detection was performed using custom scripts that automate operation of the “find by molecular feature” workflow of the Agilent Masshunter Qualitative Analysis (v7) software package. Feature alignment and recursive feature detection were performed using Agilent Mass Profiler Pro (v8.0) and Masshunter Qualitative Analysis (“find by formula” workflow), yielding an aligned table including m/z, RT, and peak areas for all features.

Data Cleaning and Degeneracy Removal: The untargeted features and named metabolites were merged to generate a combined feature table. Features missing from over 50% of all samples in a batch or over 30% of QC samples were then removed prior to downstream normalization procedure. Next, the software package Binner was utilized to remove redundancy and degeneracy in the data^98^. Briefly, Binner first performs RT-based binning, followed by clustering of features by Pearson’s correlation coefficient, and then assigns annotations for isotopes, adducts or in-source fragments by searching for known mass differences between highly correlated features.

Normalization and Quality Control: Data were then normalized using a Systematic Error Removal Using Random Forest (SERRF) approach^99^, which helps correct for drift in peak intensity over the batch using data from the QC sample runs. When necessary to correct for residual drift, peak area normalization to closest-matching internal standard was also applied to selected compounds. Both SERRF correction and internal standard normalization were implemented in R. Parameters were set to minimize batch effects and other observable drifts, as visualized using principal component analysis score plots of the full dataset. Normalization performance was also validated by examining relative standard deviation values for additional QC samples not included in the drift correction calculations.

Compound identification: Metabolites from the targeted analysis workflow were identified with high confidence (MSI level 1)^100^ by matching retention time (+/- 0.1 minute), mass (+/- 10 ppm) and isotope profile (peak height and spacing) to authentic standards. MS/MS data corresponding to unidentified features of interest from the untargeted analysis were searched against a spectral library (NIST 2020 MS/MS spectral database or other public spectral databases) to generate putative identifications (MSI level 2) or compound-class level annotations (MSI level 3) as described previously^101^.

#### LC-MS/MS untargeted lipidomics

Sample preparation: Non-targeted lipid analysis was conducted at the Georgia Institute of Technology. Powdered tissue samples (10 mg) were extracted in 400 μL isopropanol containing stable isotope-labeled internal standards (IS) with bead homogenization with 2mm zirconium oxide beads (Next Advance) using a TissueLyser II (10min, 30 Hz). Samples were then centrifuged (5 min, 21,100xg), and supernatants were transferred to autosampler vials. Plasma samples (25 μL) were extracted by mixing with 75 μL isopropanol containing the IS mix followed by centrifugation. Sample blanks, pooled extract samples used as quality controls (QC), and consortium reference samples, were prepared for analysis using the same methods. The IS mix consisted of PC (15:0-18:1(d7)), Catalog No. 791637; PE (15:0-18:1(d7)), Catalog No. 791638; PS (15:0-18:1(d7)), Catalog No. 791639; PG(15:0-18:1(d7)), Catalog No. 791640; PI(15:0-18:1(d7)), Catalog No. 791641; LPC(18:1(d7)), Catalog No. 791643; LPE(18:1(d7)); Catalog No. 791644; Chol Ester (18:1(d7)), Catalog No. 700185; DG(15:0-18:1(d7)), Catalog No. 791647; TG(15:0-18:1(d7)-15:0), Catalog No. 791648; SM(18:1(d9)), Catalog No. 791649; Cholesterol (d7), Catalog No. 700041. All internal standards were purchased from Avanti Polar Lipids (Alabaster, Alabama) and added to the extraction solvent at a final concentration in the 0.1-8 μg/ml range.

Data collection: Lipid LC-MS data were acquired using a Vanquish (ThermoFisher Scientific) chromatograph fitted with a ThermoFisher Scientific AccucoreTM C30 column (2.1 × 150 mm, 2.6 µm particle size), coupled to a high-resolution accurate mass Q-Exactive HF Orbitrap mass spectrometer (ThermoFisher Scientific) for both positive and negative ionization modes. For positive mode analysis, the mobile phases were 40:60 water:acetonitrile with 10 mM ammonium formate and 0.1% formic acid (mobile phase A), and 10:90 acetonitrile:isopropyl alcohol, with 10 mM ammonium formate and 0.1% formic acid (mobile phase B). For negative mode analysis, the mobile phases were 40:60 water:acetonitrile with 10 mM ammonium acetate (mobile phase A), and 10:90 acetonitrile:isopropyl alcohol, with 10 mM ammonium acetate (mobile phase B). The chromatographic method used for both ionization modes was the following gradient program: 0 minutes 80% A; 1 minute 40% A; 5 minutes 30% A; 5.5 minutes 15% A; 8 minutes 10% A; held 8.2 minutes to 10.5 minutes 0% A; 10.7 minutes 80% A; and held until 12 minutes. The flow rate was set at 0.40 ml/min. The column temperature was set to 50°C, and the injection volume was 2 μL.

For analysis of the organic phase the electrospray ionization source was operated at a vaporizer temperature of 425°C, a spray voltage of 3.0 kV for positive ionization mode and 2.8 kV for negative ionization mode, sheath, auxiliary, and sweep gas flows of 60, 18, and 4 (arbitrary units), respectively, and capillary temperature of 275°C. The instrument acquired full MS data with 240,000 resolution over the 150-2000 m/z range. LC-MS/MS experiments were acquired using a DDA strategy. MS2 spectra were collected with a resolution of 120,000 and the dd-MS2 were collected at a resolution of 30,000 and an isolation window of 0.4 m/z with a loop count of top 7. Stepped normalized collision energies of 10%, 30%, and 50% fragmented selected precursors in the collision cell. Dynamic exclusion was set at 7 seconds and ions with charges greater than 2 were omitted.

Data processing: Data processing steps included peak detection, spectral alignment, grouping of isotopic peaks and adduct ions, drift correction, and gap filling. Compound Discoverer V3.3 (ThermoFisher Scientific) was used to process the raw LC-MS data. Drift correction was performed on each individual feature, where a Systematic Error Removal using Random Forest (SERRF) method builds a model using the pooled QC sample peak areas across the batch and was then used to correct the peak area for that specific feature in the samples. Detected features were filtered with background and QC filters. Features with abundance lower than 5x the background signal in the sample blanks and that were not present in at least 50% of the QC pooled injections with a coefficient of variance (CV) lower than 80% (not drift corrected) and 50% (drift corrected) were removed from the dataset. Lipid annotations were accomplished based on accurate mass and relative isotopic abundances (to assign elemental formula), retention time (to assign lipid class), and MS2 fragmentation pattern matching to local spectral databases built from curated experimental data. Lipid nomenclature followed that described previously^102^.

Quality control procedures: System suitability was assessed prior to the analysis of each batch. A performance baseline for a clean instrument was established before any experiments were conducted. The mass spectrometers were mass calibrated, mass accuracy and mass resolution were checked to be within manufacturer specifications, and signal-to-noise ratios for the suite of IS checked to be at least 75% of the clean baseline values. For LC-MS assays, an IS mix consisting of 12 standards was injected to establish baseline separation parameters for each new column. The performance of the LC gradient was assessed by inspection of the column back pressure trace, which had to be stable within an acceptable range (less than 30% change). Each IS mix component was visually evaluated for chromatographic peak shape, retention time (lower than 0.2 minute drift from baseline values) and FWHM lower than 125% of the baseline measurements. The CV of the average signal intensity and CV of the IS (<=15%) in pooled samples were also checked. These pooled QC samples were used to correct for instrument sensitivity drift over the various batches using a procedure similar to that described by the Human Serum Metabolome (HUSERMET) Consortium. To evaluate the quality of the data for the samples themselves, the IS signals across the batch were monitored, PCA modeling for all samples and features before and after drift correction was conducted, and Pearson correlations calculated between each sample and the median of the QC samples.

#### Metabolomics/Lipidomics Data filtering and normalization

The untargeted metabolomics datasets were categorized as either “named”, for chemical compounds confidently identified, or “unnamed”, for compounds with specific chemical properties but without a standard chemical name. While the preprocessing steps were performed on the named and unnamed portions together, only the named portions were utilized for differential analysis. For each dataset i.e. each assay for each tissue type, the following steps are performed:

Average rows that have the same metabolite ID.

Merge the “named” and “unnamed” subparts of the untargeted datasets.

Convert negative and zero values to NAs.

Remove features with > 20% missing values.

Features with < 20% missing values are imputed either using K-Nearest Neighbor imputation (for datasets with > 12 features) or half-minimum imputation (for datasets with < 12 features).

All data are log2-transformed, and the untargeted data are median-MAD normalized if neither sample medians nor upper quartiles were significantly associated with sex or sex-stratified training group (Kruskal-Wallis p-value < 0.01). Note that for all log2 calculations, 1 is added to each value before log-normalization. This allows metabolite values that fall between 0 and 1 to have a positive log2 value.

Outlier detection was performed by examining the boxplot of each Principal Component and extending its whiskers to the predefined multiplier above and below the interquartile range (5x the IQR). Samples outside this range are flagged. All outliers were reviewed by Metabolomics CAS, and only confirmed technical outliers were removed.

#### Redundant Metabolite/Lipid Management

To address metabolites measured on multiple platforms (e.g. a metabolite measured on the HILIC positive and RP positive platforms), and metabolites with the same corresponding RefMet ID (e.g. alpha-Aminoadipic-acid, Aminoadipic acid both correspond to RefMet name ‘Aminoadipic acid’), we utilize the set of internal standards described above to make a decision on which platform’s measurement of a given feature to include in further analysis. For each tissue, based on whichever platform has the lowest coefficient of variation across all included reference standards for a given refmet id, that metabolite was chosen. The other platforms or metabolites, for this tissue, corresponding to this refmet id were removed from further downstream analysis. Data for all metabolites removed, including information about the coefficient of variation in the standards, normalized data, or differential analysis results, is available in the R Package MotrpacHumanPreSuspensionData, but is not loaded by default.

### Statistical analysis

#### Differential analysis

To model the effects of both exercise modality and time, relative to non exercising control, each measured molecular feature was treated as an outcome in a linear mixed effects model accounting for fixed effects of exercise group (RE, EE, or CON), timepoint, as well as demographic and technical covariates (see covariate selection below for more info). Participant identification was treated as a random effect. For each molecular feature, a cell-means model is fit to estimate the mean of each exercise group-timepoint combination, and all hypothesis tests are done comparing the means of the fixed effect group-timepoint combinations. In order to model the effects of exercise against non-exercising control, a difference-in-changes model was used which compared the change from pre-exercise to a during or post-exercise timepoint in one of the two exercise groups to the same change in control. This effect is sometimes referred to as a “delta-delta” or “difference-in-differences” model.

#### Model selection via simulation

In order to determine the optimal statistical package/model for these data – given MoTrPAC’s unique sampling design (see ‘acute exercise intervention’ above) – a simulation study was conducted to determine the impact of various approaches to account for correlation in repeated measurements. The simulation evaluated type 1 error, power, and bias for multiple plausible modeling approaches, to identify those that would obtain higher power while maintaining nominal type 1 error rates. For a subset of molecular features in the datasets, mean, covariance, skew, and kurtosis were summarized over the relevant timepoints in the acute exercise bout within subgroups defined by sex and exercise type. Then, these summary values were used in combination with the “covsim” R package to generate non-normally distributed simulated data^103^. Additionally, sample size and missingness patterns were aligned in the simulated data to match those of the observed data. Using a total of 5 million simulated instances where data were generated under the null hypothesis (i.e., no change in the mean values of the outcome over the exercise timepoints) or alternative hypothesis (i.e., the mean value of the outcome differs between at least two timepoints), type 1 error rate and power, respectively, were evaluated for seven analytic strategies: (1) pairwise t-tests between timepoints of interest, (2) ordinary least squares regression, (3) differential expression for repeated measures (i.e., dream) with random intercepts,129 (4) linear mixed models with random intercepts, (5) mixed models for repeated measures using an unstructured covariance matrix, and (6) generalized linear models using generalized estimating equations (GEEs) and a first degree autoregressive correlation structure. These analyses were replicated for adipose, blood, and muscle samples. The dream function obtained type 1 error rates 5.2, 5.4, and 5.2% for adipose, blood, and muscle analyses, respectively, with higher power than all other methods apart from generalized linear models with GEEs. Although generalized linear models with GEEs had higher power than dream, they also had higher type 1 error with 7.7%, 6.0%, and 6.5% type 1 error rates for adipose, blood, and muscle analyses, respectively. Therefore, due to stability of type 1 error and relatively high power compared to other approaches, the dream function from the variancePartition R package was selected for the primary analyses for every omic platform except Methyl-Cap (see methylation capture sequencing methods above for more details).129

#### Covariate selection

Covariates were selected through a combination of a priori knowledge about factors influencing molecular levels as well through empiric screening. Factors were considered for model inclusion by correlating the principal components of each tissue-ome feature set to demographic and technical factors using variancePartition::canCorPairs^78^. Visualizations and computational analysis that describe this process for the decisions for covariate selection can be found at: https://github.com/MoTrPAC/precovid-analyses/tree/main/QC. Ultimately, the following fixed effect covariates were included in the models for every omic platform, except Methyl Cap (see methods above): group and timepoint in a cell means model, clinical site, age, sex, and BMI. ‘Participant id’ was included as a random effect in every model. Each ome then had ome-specific covariates selected as described in prior sections.

A targeted analysis was conducted to assess whether and how race, ethnicity, and genetic ancestry could be incorporated, given prior evidence that these factors can influence omic data^104,105^. We evaluated multiple models that included patient-reported race and/or ethnicity, as well as principal components derived from whole-genome sequencing (WGS), to determine their impact on model fit using the BIC. However, no combinations of individual or joint inclusion of race, ethnicity, or WGS-derived principal components led to an improvement in average model BIC across all features for a platform.

The final set of covariates included in each model for each tissue and ome can be found in the R Package MotrpacHumanPreSuspensionData.

#### Missingness

Data missingness varied for multiple reasons including experimental design (i.e. temporal randomization which randomized some participants to have samples obtained at a subset of timepoints. For some omic sets, imputation to alleviate missingness could be performed (metabolomics) but for others was not, including proteomics. Analysis demonstrated proteomics missingness was approximately at random (data not shown).

Given the above issues affecting the ultimate sample size for effect estimation, for a feature to be included in the analysis, a minimum number of 3 participants were required to have a paired pre-exercise sample for all groups and all during/post-exercise timepoints. Thus, all group comparisons (e.g. RE vs CON at 3.5 hours post-exercise) required 3 participants with matched pre- and post-exercise samples in each group. This requirement was satisfied in all omes and tissues except a subset of MS-acquired proteomics and phosphoproteomics features. Features not meeting this criterion in any group-timepoint set were excluded from differential analysis entirely, though raw and QC-normalized values are available.

#### Specific omic-level statistical considerations

Models for all omes other than methylation were fit using ‘variancePartition::dream’^78^. For the transcriptomics and ATAC-seq, which are measured as numbers of counts, the mean-variance relationship was measured using ‘variancePartition::voomWithDreamWeights’ as previously described^78^. For all other omes the normalized values were used directly as input to the statistical model.

The full implementation of the statistical models for each of non-methylation datasets can be found in the R Package MotrpacHumanPreSuspensionData. Methylation data was processed separately, and the methods can be found in the methylation methods section.

#### Contrast types

The specific contrasts made in the statistical analysis include three categories of comparison: difference-in-changes relative to control, group-specific, and resistance vs endurance. All comparisons were structured using ‘variancePartition::makeContrastsDream’^78^.

The difference-in-changes model as described at the opening of this section represents the primary DA analysis, and any individual feature mentioned as statistically significantly changing due to exercise will be referring to the difference-in-changes results unless otherwise specifically stated.

The next category of contrasts is a group-specific comparison, which compares a given post-intervention timepoint to pre-exercise within a given group, without comparing to the control group. This contrast was implemented to more easily quantify effect sizes within each group independently, but are not used as a general endpoint.

Finally, the RE vs EE is very similar to the difference-in-changes model, but sets the RE group as a matched control to the EE group. In this comparison, the change in response to exercise at a given timepoint in one exercise group is directly compared to the corresponding change in the other exercise group. This contrast can directly test the hypothesis that the EE effect is different from the RE effect. There are no samples obtained during RE bouts, so those timepoints in blood were excluded from this comparison.

#### Significance thresholds

For each of the above contrasts, p-values were adjusted for multiple comparisons for each unique contrast-group-tissue-ome-timepoint combination separately using the Benjamini-Hochberg method to control False Discovery Rate (FDR)^106^. Features were considered significant at a FDR of 0.05 unless otherwise stated.

#### Human feature to gene mapping

The feature-to-gene map links each feature tested in differential analysis to a gene, using Ensembl version 105 (mapped to GENCODE 39)^107^ as the gene identifier source. Proteomics feature IDs (UniProt IDs) were mapped to gene symbols and Entrez IDs using UniProt’s mapping files^108^. Epigenomics features were mapped to the nearest gene using the ChIPseeker::annotatePeak()^109,110^ function with Homo sapiens Ensembl release 105 gene annotations. Gene symbols, Entrez IDs, and Ensembl IDs were assigned to features using biomaRt version 2.58.2 (Bioconductor 3.18)^111–113^.

Metabolite features were mapped to KEGG IDs using KEGGREST, RefMet REST API, or web scraping from the Metabolomics Workbench^112,114^.

### Enrichment analysis

#### Preparation of differential analysis results

The differential analysis results tables for each combination of tissue and ome were converted to matrices of z-scores with either gene symbols, RefMet metabolite/lipid IDs, or phosphorylation flanking sequences as rows and contrasts as columns. These matrices serve as input for the enrichment analyses. For the transcriptomics, TMT proteomics, and Olink proteomics results, transcripts and proteins were first mapped to gene symbols. To resolve cases where multiple features mapped to a single gene, only the most extreme z-score for each combination of tissue, contrast, and gene was retained. Metabolite and lipid identifiers were standardized using the Metabolomic Workbench Reference List of Metabolite Names (RefMet) database^115^. For the CAMERA-PR enrichment of phosphoproteomics results, since peptides could be phosphorylated at multiple positions, phosphorylation sites were separated into single sites with identical row information (e.g., protein_S1;S2 becomes protein_S1 and protein_S2, both with the same data). For any combinations of contrast and phosphosite that were not uniquely defined, only the most extreme z-score (maximum absolute value) was selected for inclusion in the matrix. Singly phosphorylated sites were required to use the curated kinase–substrate relationship information available from PhosphoSitePlus (PSP)^108^, described below.

#### Gene set selection

Gene sets were obtained from the MitoCarta3.0 database, CellMarker 2.0 database^116^, and the C2-CP (excluding KEGG_LEGACY) and C5 collections of the human Molecular Signatures Database (MSigDB; v2023.2.Hs)^116–118^ Metabolites and lipids were grouped according to chemical subclasses from the RefMet database^115^. This includes subclasses such as “Acyl carnitines” and “Saturated fatty acids”.Flanking sequences for protein phosphorylation sites were grouped according to their known human protein kinases provided in the Kinase_Substrate_Dataset file from PhosphoSitePlus (PSP; v6.7.1.1; https://www.phosphosite.org/staticDownloads.action; last modified 2023-11-17)^108^. In addition to these kinase sets, sets from the directional Post-Translational Modification Signatures Database (PTMsigDB) were included for analysis^119^.

For each combination of tissue and ome, molecular signatures were filtered to only those genes, metabolites/lipids, or flanking sequences that appeared in the differential analysis results. After filtering, all molecular signatures were required to contain at least 5 features, with no restriction on the maximum size of sets. Additionally, gene sets were only kept if they retained at least 70% of their original genes to increase the likelihood that the genes that remain in a given set are accurately described by the set label.

#### Analysis of Molecular Signatures

Analysis of molecular signatures was carried out with the pre-ranked Correlation Adjusted MEan RAnk (CAMERA-PR) gene set test using the z-score matrices described in the “Preparation of differential analysis results” section to summarize the differential analysis results for each contrast at the level of molecular signatures^25,119^. Z-scores were selected as the input statistics primarily to satisfy the normality assumption of CAMERA-PR, and the analysis was carried out with the cameraPR.matrix function from the TMSig R/Bioconductor package^120,121^.

#### Over-representation analysis (ORA)

Over-representation analysis (ORA) was conducted using the run_ORA function from the MotrpacHumanPreSuspension R package (v0.0.1.53) which uses the hypergeometric test for statistical significance. The input consisted of differentially expressed genes (adjusted p-value < 0.05) identified from one or more omic layers in a particular tissue. The background gene set included all genes detected in each of the ome(s) under consideration.

### Statistical significance thresholds

For both CAMERA-PR and ORA results, p-values were adjusted within each combination of tissue, ome, contrast, and broad molecular signature collection (MitoCarta3.0, CellMarker 2.0, C2, C5, RefMet, PSP, and PTMsigDB) using the Bejamini-Hochberg method. Gene sets and RefMet subclasses were declared significant if their adjusted p-values were less than 0.05. This threshold was raised to 0.1 for kinase sets.

#### Visualization methods

Bubble heatmaps were generated from the enrichment analysis results using the enrichmap function from the TMSig R/Bioconductor package^120^.

#### Clustering techniques

##### Fuzzy C-Means Clustering

The z-score matrices described in the Enrichment Analysis Methods were used as input for fuzzy c-means (FCM) clustering^122,123^. The matrices for the different omes, excluding global proteomics and phosphoproteomics (as these were only measured at two timepoints) were stacked to form a single matrix with tissue-specific timepoints as columns and features being a row. The rows of this matrix were then divided by their sample standard deviations, which were calculated after two columns of zeros were temporarily included to represent the pre-exercise timepoints; these zero columns were discarded before clustering. The Mfuzz R package was used to perform FCM clustering with the parameter m set to the value of Mfuzz::mestimate^124,125^. The minimum distance between centroids was used as the cluster validity index to determine the optimal number of clusters for each tissue. A scree/elbow plot of these distances for a range of cluster numbers (3-14) was generated for each tissue, and the final cluster numbers were determined through manual inspection (n=13). For visualization purposes, hard clustering was performed by assigning each feature to the cluster with the highest membership probability; features with probabilities below 0.3 are not clustered or displayed. Cluster numbers are assigned arbitrarily.

### Sex difference

To compare the differential abundance of features between male and female participants, we fit a linear model with a single covariate (participant sex). Several alternative models were fit for models adjusting the effect of age, clinical site, technical covariates, etc. on molecular expression. Each of these models had, on average across all features for a given platform, a weaker model fit as determined by Bayesian Information Criteria (BIC), partially due to the limited number of samples available to estimate these additional parameters. The results and model fits used to guide this decision are available in: https://github.com/MoTrPAC/precovid-analyses/blob/main/figures/adipose/baseline_sex_differences.Rmd

### Cell type analysis

#### Cell type enrichment

ASAT cell marker genes were curated from an integrated object of two previously published ASAT full-length single-nuclei (sn) RNAseq^21,40^. FindALLMarkers() from Seurat R package was used to identify marker genes (min.pct=0.1, logfc.threshold=0.25), and adjusted p-value<0.05 and average log2FC>1 were further applied to filter out non-significant and non-specific marker genes. A total of nine cell types were included: two mature adipocyte subtypes (i.e., Adipocyte-1 and -2), pre-adipocyte, stem cell, vascular cell, two macrophage subtypes (i.e., M1-like lipid-associated macrophage, LAM and M2-like resident macrophage), mast cell, and NK/T cell. Each cell type consisted of a list of 100 marker genes (Table S2). CAMERA-PR was applied on ASAT transcriptomics data using curated cell-type marker genes as a reference geneset.

#### Deconvolution of ASAT cell types

Deconvolution—i.e., in silico estimation of cell type proportions—was performed on ASAT transcriptomic data as previously described^22^. Briefly, an ASAT-specific cell type signature matrix was derived from full-length single-nucleus RNA-seq data generated from ASAT of 20 adults^21^. Using the optimized pipeline for human ASAT deconvolution^22^, we estimated the relative proportions of seven distinct ASAT cell populations: two mature adipocyte subtypes (Adipocyte-1, Adipocyte-2), vascular cells, stem cells, pre-adipocytes, macrophages, and mast cells. Estimated proportions were computed across all bulk RNA-seq samples using this reference signature.

#### Weighted Gene Co-expression Network Analysis (WGCNA)

WGCNA (version 1.73) was employed to identify distinct, non-overlapping clusters (modules) in each ASAT omic layer at baseline samples (transcriptomics, n=172; proteomics and phosphoproteomics, n=42; metabolomics, n=172). Connectivity for each omic feature was calculated by summing its correlation strengths with all other features. A scale-free signed network was constructed using an optimal soft-threshold power (β>0.8). A dynamic tree-cutting algorithm and a merging threshold of 0.3 was used for all omic layers.

In each omic layer, module 0 (i.e., T0, Pr0, Ph0, and M0) often referred to as the ‘grey’ module, comprises transcripts that could not be assigned to any other module due to weak correlation patterns. Therefore, module 0 was excluded for downstream analyses. Top hub genes within each module were determined based on their kME values (module eigengene-based connectivity). Module eigengenes (MEs), representing the first principal eigenvector of each module, were extracted and correlated with clinical and subclinical variables using biweight midcorrelation. ORA was conducted on module features. For phosphoproteomics, phosphosites were collapsed and the original protein was used. For metabolomics, REFMET was used as a reference database.

### Secretome analysis

A list of putative secreted proteins was curated by combining two sources: proteins detected in plasma via Olink proteomics (n = 1,408) (*REF: MoTrPAC Blood Companion, in review*) and the known secretome annotated in the Human Protein Atlas (n = 2,520)^51^, resulting in 3,277 unique proteins. Any differentially regulated transcripts or proteins in ASAT that overlapped with this curated list were considered candidate *exerkines*—putative adipose-derived secretory factors responsive to acute exercise.

Extracellular score (“COMPARTMENTS”). Subcellular localization of proteins was annotated using the COMPARTMENTS database^53^, which provides scores ranging from 0 to 5, with higher scores indicating greater confidence in localization. The “All channels integrated” file for human proteins was downloaded (https://download.jensenlab.org/human_compartment_integrated_full.tsv). To calculate the extracellular score, we selected the highest score among the following compartments associated with extracellular localization: *Extracellular region*, *Extracellular space*, *Extracellular exosome*, and *Extracellular vesicle*.

#### Gene Derived Correlation Across Tissues (GD-CAT)

Transcriptomic data from the GTEx (Genotype-Tissue Expression) project (nL=L310 individuals, 19 tissues) were accessed via the GD-CAT pipeline^52,54^. GD-CAT computes gene-to-gene correlations between user-defined origin and target tissues to infer potential inter-tissue regulatory relationships. Of the 60 ASAT-exerkine candidate genes identified in our study, 48 were detected in the subcutaneous adipose tissue (SAT) transcriptome of the GTEx cohort. These 48 SAT genes were correlated with the expression of all genes in 18 distal target tissues. Correlation was performed using biweight midcorrelation (bicor), and significance was determined at adjusted p < 0.05 following Benjamini-Hochberg correction for multiple testing. To assess the biological pathways associated with SAT-derived exerkine signaling, bicor values were z-score transformed, and CAMERA-PR was applied to the z-scaled bicor vectors of correlated genes in each distal tissue.

### Integration of MoTrPAC rat endurance training study

Quantitative Endocrine Network Interaction Estimation (QENIE) quantifies the potential endocrine influence of secretory genes by calculating a secretome score (Ssec) for each gene in a given origin tissue, based on gene–gene correlation with target tissues^59^. The pipeline begins by computing a p-value matrix from biweight midcorrelations between gene expression in the origin and target tissues. For each secretory gene in the origin tissue, the sum of -ln(p-value) across all genes in the target tissue is computed as the raw Ssec. This score is then normalized by the number of genes in the target tissue, allowing for comparisons across origin–target tissue pairs. Higher Ssec values indicate a stronger overall correlation—and potential endocrine influence—between a secretory gene in the origin tissue and transcriptional programs in a given target tissue. QENIE was applied to multi-tissue transcriptomic datasets from the MoTrPAC rat endurance training study as previously described^58^. Briefly, subcutaneous white adipose tissue (WATSC) was designated as the origin tissue. Sex-adjusted correlation p-value matrices were obtained from correlating WATSC transcript expression matrix against 16 other tissues within each training group (i.e., TR1W, 1-week training; TR2W, 2-week training; TR4W, 4-week training; TR8W, 8-week training, and a non-exercising control group (CON) duration-matched with TR8W; n=10 (5F, 5M) for each group). Ovary, testis (due to differing sample sizes), and brown adipose tissue (due to its limited presence in adult humans) were excluded. Subsequently, only secretory protein-coding genes were retained using the same curated list described above, yielding a total of 2,248 secretory genes identified in WATSC for Ssec calculation. Density plots of Ssec distributions were generated across all WATSC-to–target tissue pairs. Wilcoxon signed-rank tests and paired t-tests were used to compare Ssec ranks between CON and TR8W for the 53 mappable ASAT-exerkine candidate genes. As previously described, fast gene set enrichment analysis (FGSEA) was performed on bicor vectors of genes in vastus lateralis and liver that correlated with WATSC-IGFBP7 at each time point to assess functional associations.

## QUANTIFICATION AND STATISTICAL ANALYSIS

Statistical approaches for differential and enrichment analyses are described in their respective Methods sections. Biweight midcorrelation was performed using WGCNA::bicor() for Figures 1C, S1A, 5B–D, S5A–C, 6D, S6B–C, as well as for the secretory score analyses (i.e., QENIE and GD-CAT) in Figures 7 and S7. Spearman correlation was applied for Figure S2I. Student’s *t*-test (independent) was used for Figure S1B, and the paired *t*-test was used for Figure S7A. A two-way linear mixed-effects model was conducted for Figure S4A using lme4::lmer(), with exercise group and time point as fixed effects. Adjusted *p*-values were calculated using the Benjamini–Hochberg method. All statistical analyses were performed in R (R Foundation for Statistical Computing, Vienna, Austria)

---

**Figure S1.**
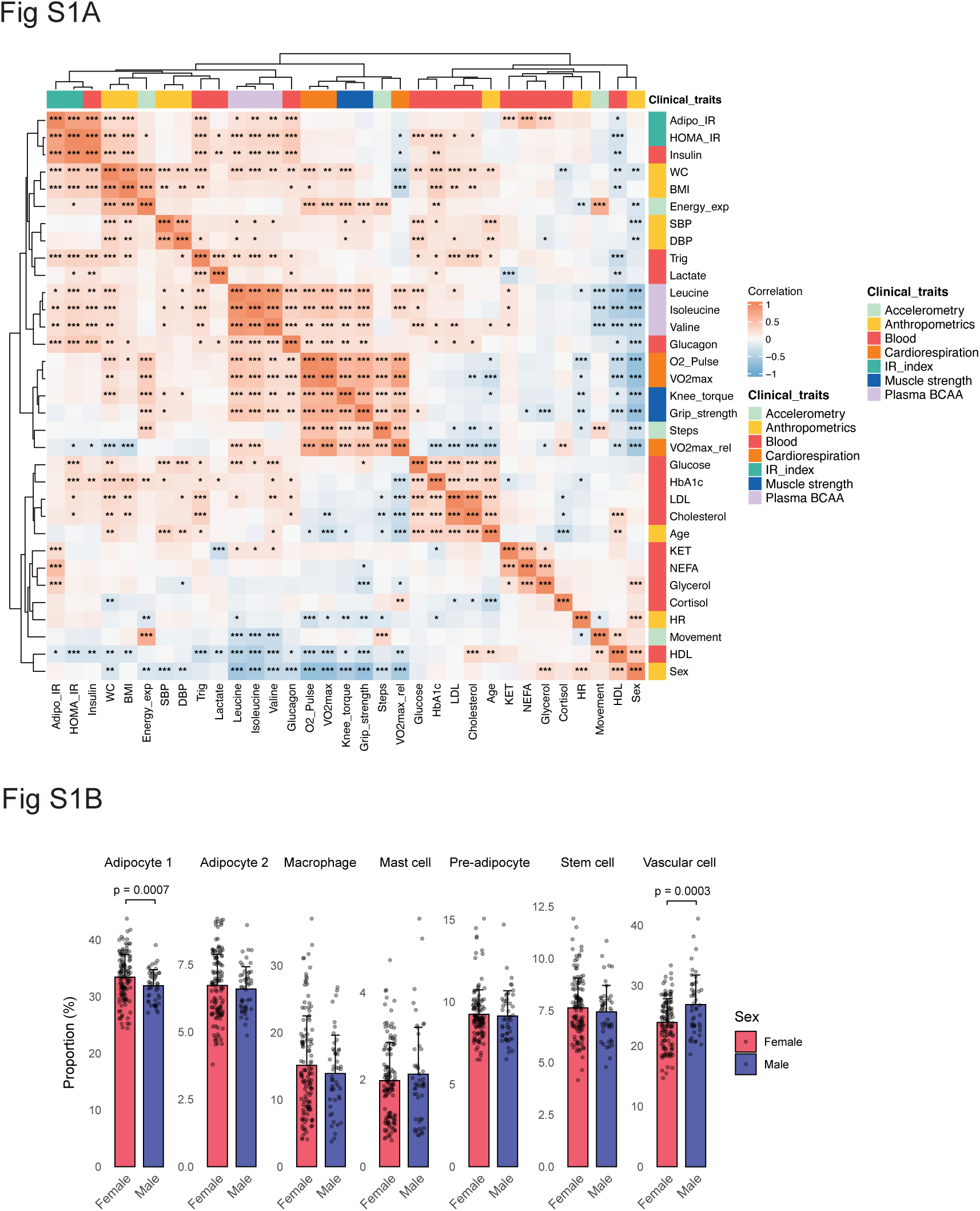
Intercorrelation of clinical/sub-clinical variables and sexual dimorphism in ASAT cell type proportions at baseline (Supplement to Figure 1) A) Intercorrelation heatmap across clinical and subclinical variables. “p<0.05, **p<0.01, ***p<0.001. B) Sex-based comparisons of deconvoluted ASAT cell type proportions. BMI, Body mass index; WC, Waist circumference; SBP, Systolic blood pressure; DBP, DIastolic blood pressure; HR, Heart rate; NEFA, Non-esterifed fatty acid; Total_C, Total cholesterol; HDL, High-density lipoprotein; LDL, Low-density lipoprotein.

**Figure S2.**
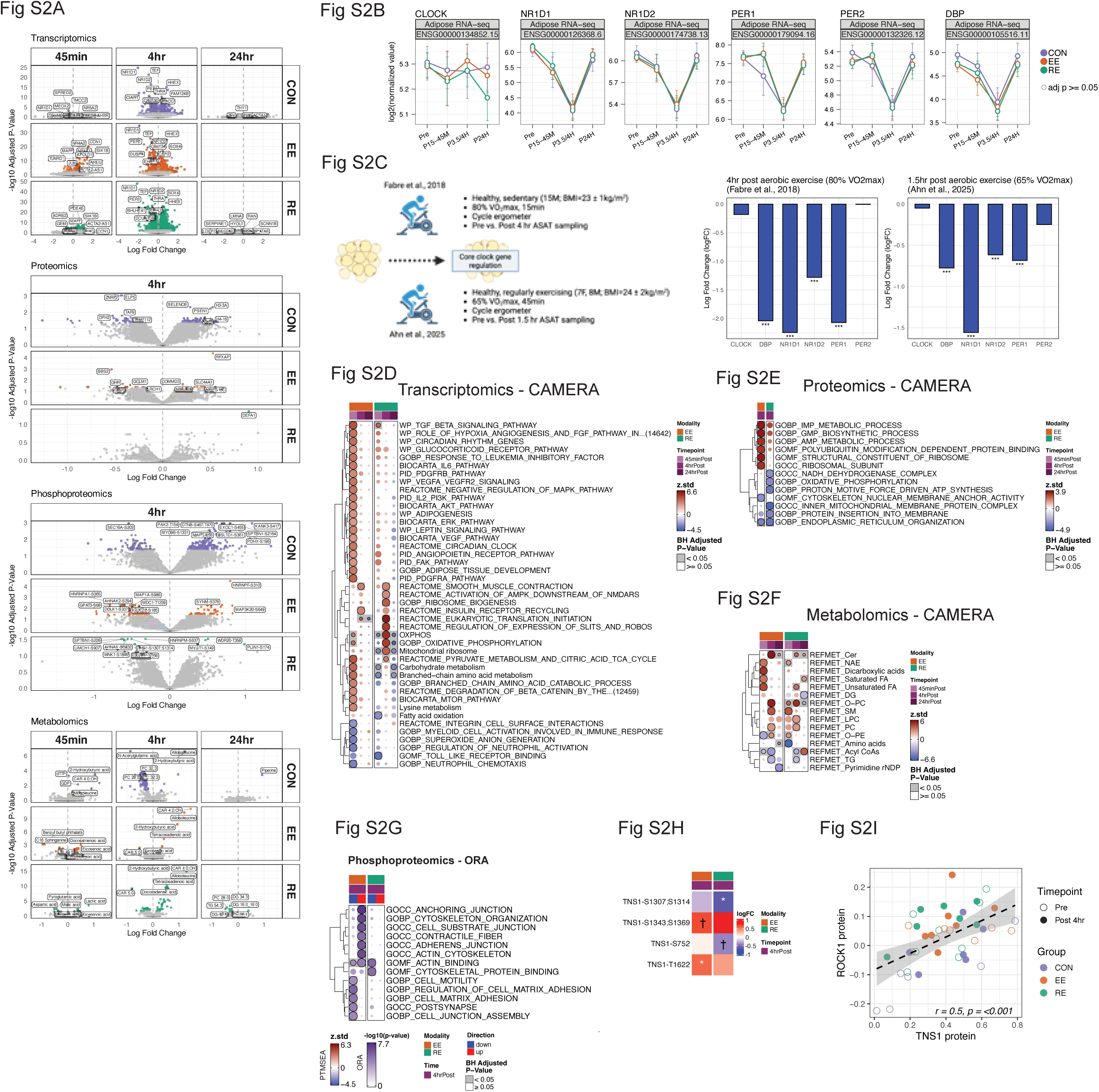
Differential analysis and enrichment analysis (Supplement to Figure 2) A) volcano plots showing differentially regulated features across omic layers and timepoints in CON, and in CON-unadjusted EE and RE groups. B) Temporal expression patterns of core circadian transcripts (CLOCK, NR1D1/2, and PER1/2, and DBP) in response to exercise. C) Acute exercise effects on core clock genes from previous reports (Fabre et al., 2018 and Ahn et al., 2025). In Fabre et al., 15 sedentary males without obesity performed 15min of moderate intensity endurance exercise (80% VO_2_ max) on a cycle ergometer. ASAT sampling occurred 4 hr post-exercise. In Ahn et al., 15 regular exercisers (7 females and 8 males) without obesity Performed 45min of moderate intensity endurance exercise (65% VO_2_ max) on a cycle ergometer. ASAT sampling occurred 1.5 hr post-exercise. In both studies, ASAT transcriptome was profiles via bulk RNA sequencing. Bar plots show changes in core clock genes post-exercise in two studies. *adjusted p<0.05, **adjusted p<0.01, ***adjusted p<0.001. D) CAMERA-PR analysis of transcriptomic, E) proteomic, and F) metabolomic changes post-exercise. G) ORA results on collapsed differentially regulated phosphoprotein, stratified by directionality. H) Heatmap illustrating exercise responses on TNS1 phosphorylation at post 4 hr. *adjusted p<0.05, †adjusted p<0.1. I) Spearman correlation between protein abundance of ROCK1 and TNS1.

**Figure S3.**
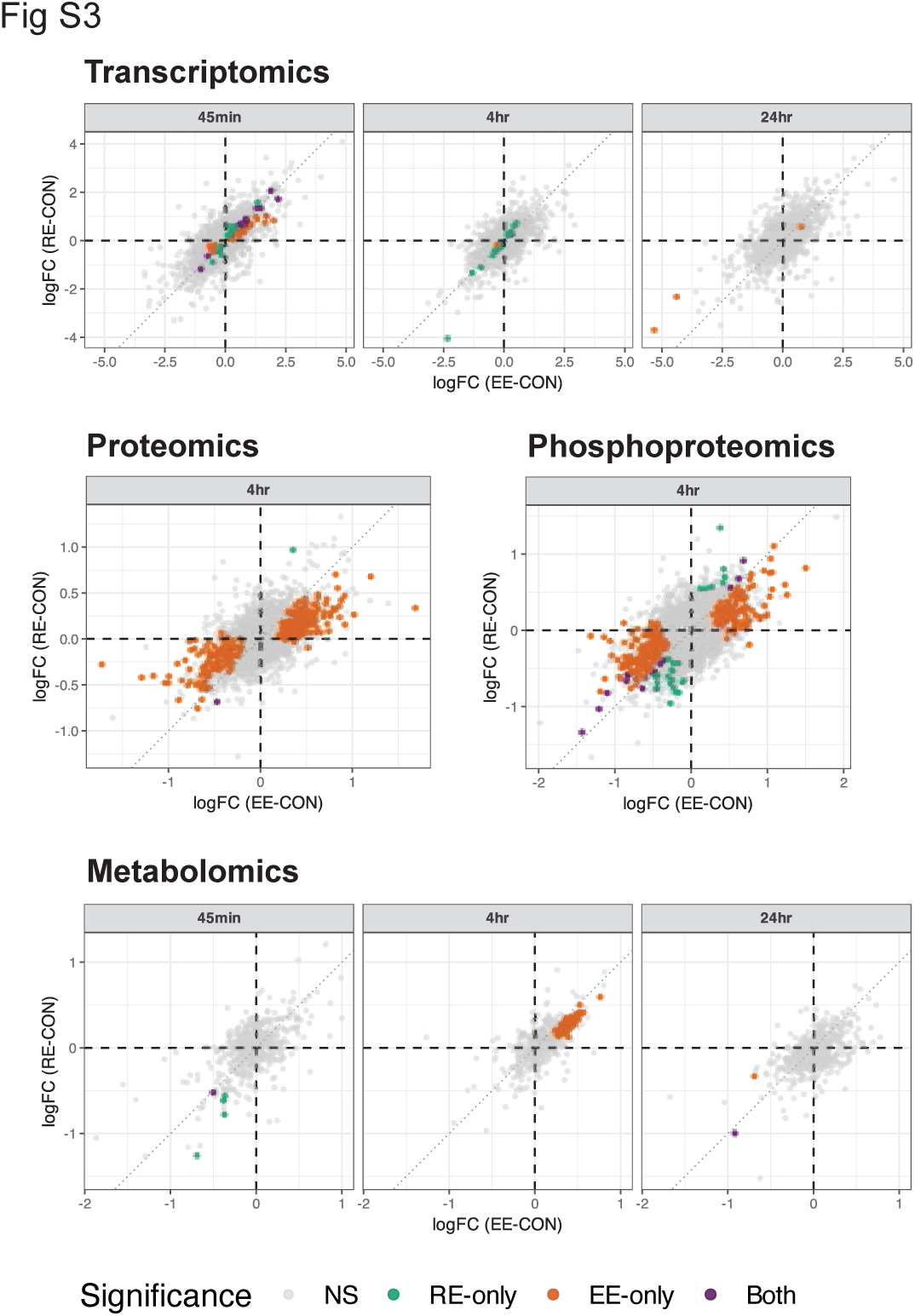
Comparison of EE vs. RE (Supplementary to Figure 3) Scatter plots comparing logFC between EE and RE for features differentially regulated (adjusted p<0.05) in EE or RE.d p<0.05.

**Figure S4.**
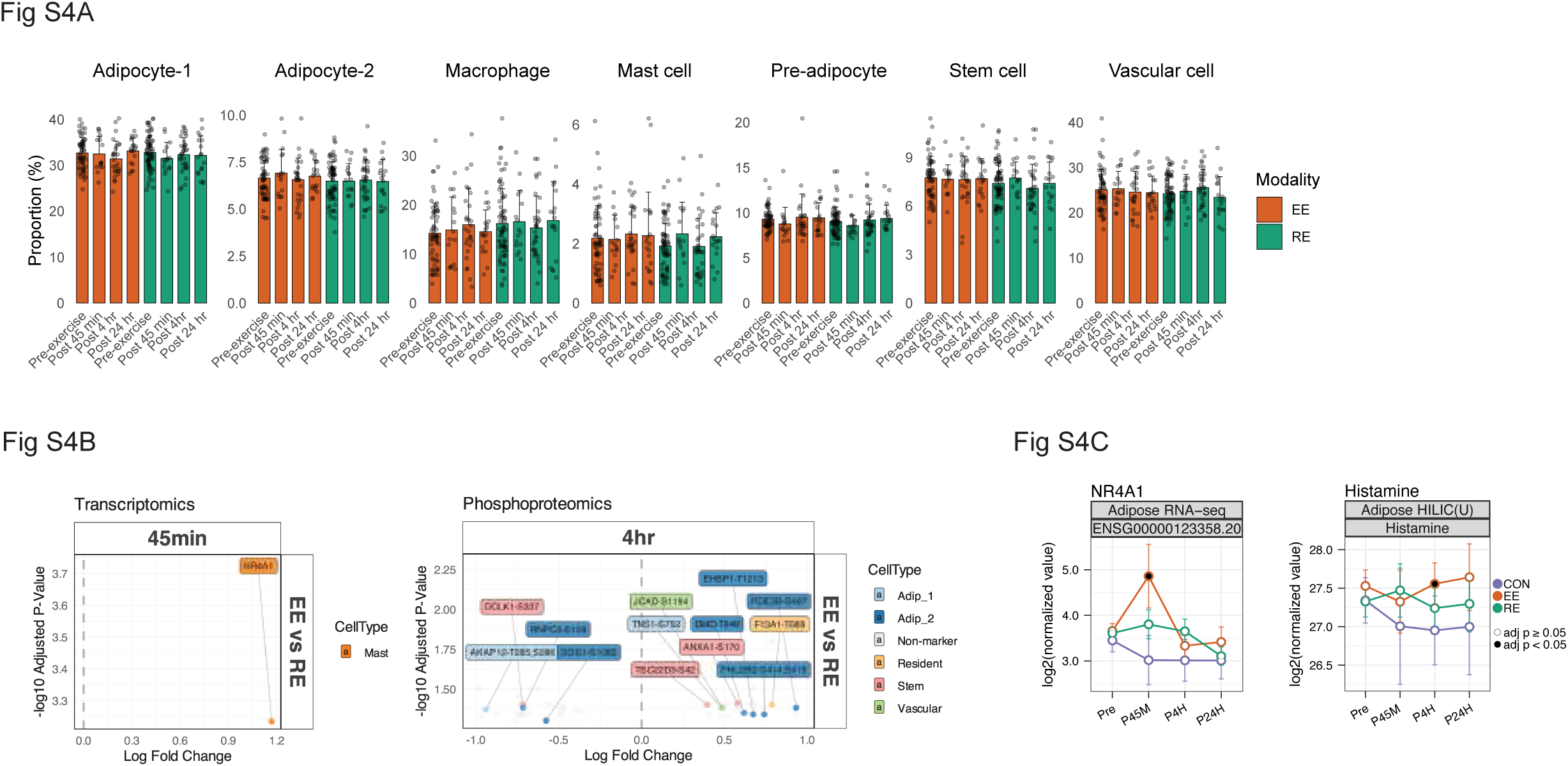
Cell type deconvolution and enrichment (Supplement to Figure 4) A) Estimated ASAT cell type proportions (%) from deconvolution analysis at each timepoint, separated by exercise modality. No significant modality × timepoint interaction or main effect was detected. B) Volcano plots of differentially regulated phosphosites in EE vs. RE, focusing on features annotated as cell-type markers. logFC > 0 indicates greater enrichment in EE; logFC < 0 indicates greater enrichment in RE. C) Temporal expression patterns of *NR4A1* and histamine in ASAT.

**Figure S5.**
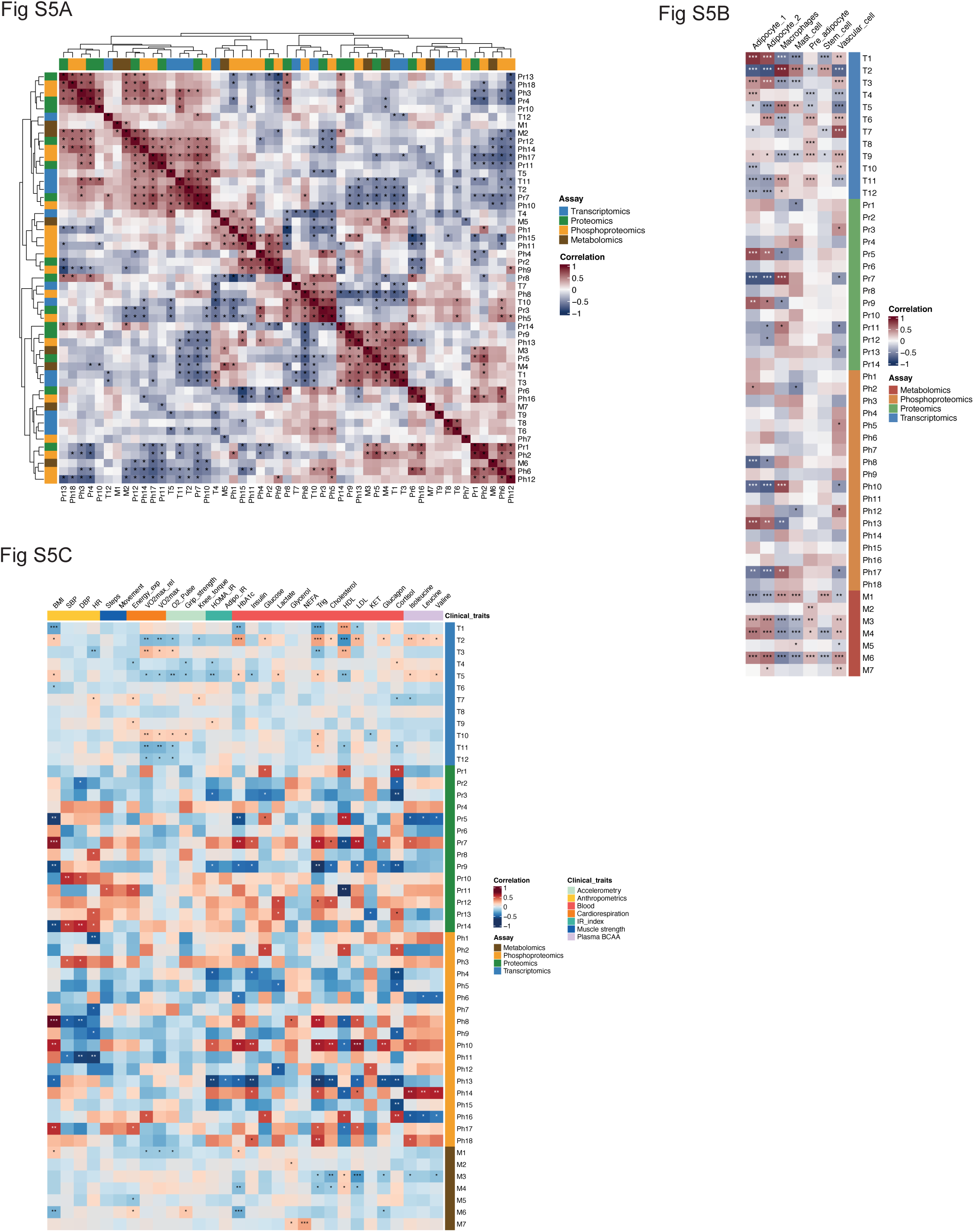
Integrative network downstream correlation analyses (Supplement to Figure 5) A) Heatmap illustrating cross-omic correlation of constructed modules. *p<0.05. B) Heatmap showing unadjusted biweight midcorrelations between module eigengenes and estimated ASAT cell type proportions from baseline deconvolution. *p<0.05; **p<0.01; ***p<0.001. C) Heatmap showing biweight midcorrelations between module eigengenes and clinical traits, adjusted for sex, age group, and waist circumference (WC). Covariate adjustment was performed by regressing both traits and eigengenes, and residuals were used for correlation. *p<0.05; **p<0.01; ***p<0.001.

**Figure S6.**
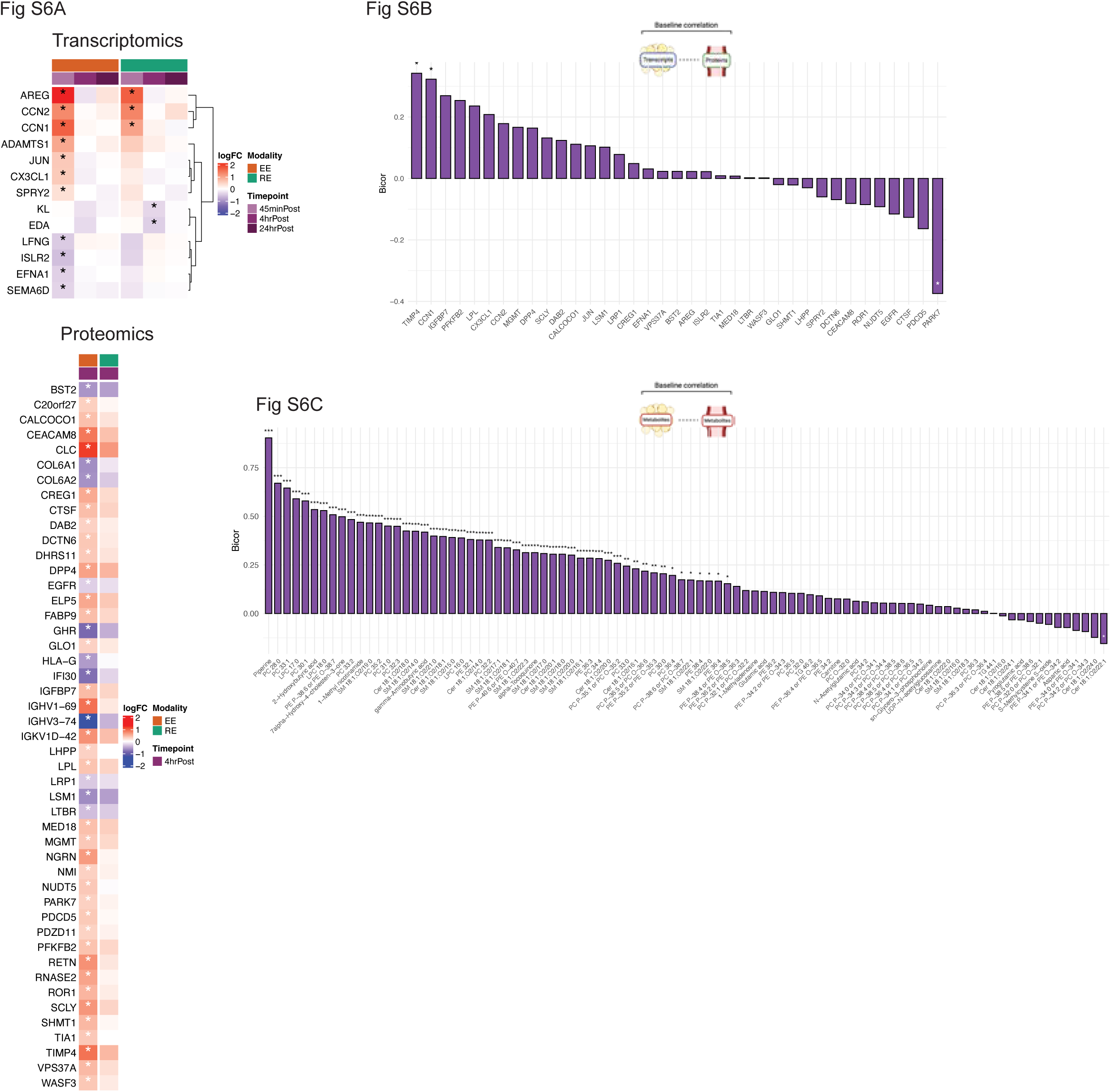
Relationship between ASAT feature abundance and corresponding plasma feature abundance at baseline (Supplementary to Figure 6) A) Heatmaps illustrating exercise responses on filtered exerkine candidates. *adjusted p<0.05. B) Biweight midcor-relation between ASAT-exerkine transcript expression and corresponding plasma protein abundance at baseline. *p<0.05. C) Biweight midcorrelation between ASAT-exerkine metabolites and corresponding plasma metabolites at baseline. *p<0.05, **p<0.01, ***p<0.001

**Figure S7.**
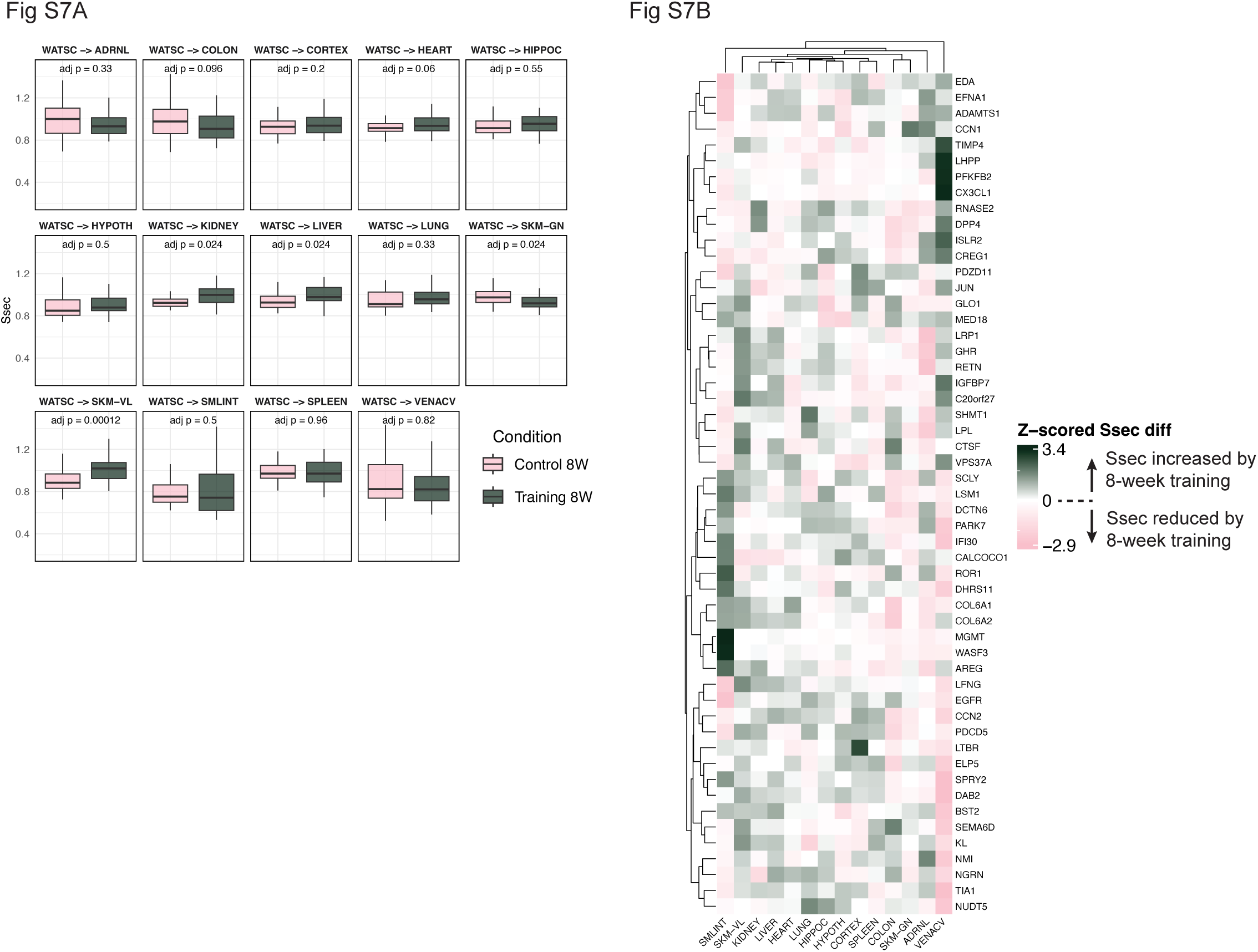
ASAT-exerkine candidates in endurance training (Supplementary to Figure 7) A) Paired T-test results comparing Ssec for each WATSC-to-target tissue connection between CON and Pink boxplot refers to CON. Dark green boxplot refers to TR8W. B) Heatmap showing Ssec difference of 60 ASAT-secreted candidates between TR8W vs. CON. Positive z-scaled Ssec diff indicates higher Ssec observed in the 8-week trained rats compared with 8-week untrained rats for the given WATSC-to-target tissue connection.

## Notes

https://motrpac-data.org/

### Competing Interest Statement

The authors have declared no competing interest.

### Summary of Updates

The author names have been corrected

